# Regulation of protein translation by TRIM21

**DOI:** 10.1101/2022.06.07.495079

**Authors:** Huiyi Li, Shun Liu, Qing Feng, Rilin Deng, Jingjing Wang, Xintao Wang, Renyun Tian, Yan Xu, Shengwen Chen, Qian Liu, Luoling Wang, Xinran Li, Mengyu Wan, Yousong Peng, Songqing Tang, Binbin Xue, Haizhen Zhu

**Affiliations:** Institute of Pathogen Biology and Immunology of College of Biology, Hunan Provincial Key Laboratory of Medical Virology, State Key Laboratory of Chemo/Biosensing and Chemometrics, Hunan University, Changsha, Hunan, China; Bioinformatics Center of College of Biology, Hunan Provincial Key Laboratory of Medical Virology, Hunan University, Changsha, Hunan, China; Research Center of Cancer Prevention and Treatment, Translational Medicine Research Center of Liver Cancer, Hunan Cancer Hospital, Changsha, Hunan, China

## Abstract

Regulation of translation initiation is essential for maintenance of protein homeostasis and typically involves the phosphorylation of translation initiation factor eIF2α in eukaryotes. In response to stressors, cells employ eIF2α-dependent signaling to control translation initiation, which regulates multiple biological and physiological processes. However, the precise regulatory mechanism remains unclear. In this study, we focus on the role of TRIM21 in the regulation of protein translation mediated by protein kinase R (PKR), one of the classic kinases that phosphorylates eIF2α. TRIM21 deficiency enhances the activation of PKR under different types of stress. TRIM21 interacts with PKR, and the E3 ligase activity of TRIM21 is crucial for stress-triggered PKR inactivation. TRIM21 interacts with the PKR phosphatase PP1α and promotes K6-linked polyubiquitination of PP1α under stresses. Ubiquitination of PP1α augments its interaction with PKR, causing PKR inactivation and subsequent dephosphorylation of eIF2α, initiating protein synthesis. Moreover, TRIM21 constitutively restricts viral infection by reversing PKR-mediated inhibition of the protein synthesis of intrinsic antiviral genes. The TRIM21-PP1α axis acts as a newly discovered program that regulates PKR-associated protein synthesis and broadens our knowledge of antiviral genes. Moreover, TRIM21-mediated regulation of PKR activation provides evidence that TRIM21 may anticipate the interferon-dependent immunotherapy. Our study highlights the essential role of TRIM21 in regulating protein translation and may provide a novel target for the treatment of translation-associated diseases.

## Introduction

Protein translation is a basic biological process among all species and includes three steps: initiation, elongation and termination. Appropriate regulation of protein translation is essential for the maintenance of protein homeostasis, and disruption of protein translation by extrinsic or intrinsic stress is responsible for severe organ damage, contributing to tumorigenesis and inflammation (Wek RC, 2018). In eukaryotes, cells commonly employ stress-induced programs, inhibiting global initiation of translation in the cell, which conserves energy and facilitates reprogramming of gene expression to restore protein homeostasis and promote adaptation to the alteration of the cellular environment (Talloczy Z et al., 2002). The central mechanism regulating translation initiation in response to stress involves the phosphorylation of translation initiation factor eIF2α (Pakos-Zebrucka K et al., 2016). There are four kinases, dsRNA-dependent protein kinase PKR, GCN2 (General Control Nonrepressed 2), PERK (PKR-like Endoplasmic Reticulum Kinase) and HRI (Heme-Regulated eIF2α Kinase), responsible for eIF2α phosphorylation in eukaryotic cells (Hotamisligil GS, 2010). Among them, as one of the pattern recognition receptors (PRRs) involved in innate immunity, PKR is distinct for its essential role in the response to viral or bacterial infection by detecting dsRNA or binding to inflammasome-associated receptors, which tightly links stress and immunity (Chen YG and Hur S, 2022). In addition to being activated by dsRNA derived from extracellular virus, PKR is also activated by endogenous dsRNAs, such as those from endogenous retroviral elements (EREs), which have been reported to be involved in interferon-dependent antitumor response (Chen R et al., 2021). Several lines of evidence indicate that the dsRNA derived from EREs benefits for immunotherapy, which can trigger PKR activation, contributing to protein translational shutdown and apoptosis in cancer cells (Chen R et al., 2021, Zhang T et al., 2022). However, how host regulates PKR activation remains unclear.

Environmental stress, such as viral infection or double-stranded RNA (dsRNA) replication intermediates generated in virus-infected cells, triggers a PKR-dependent stress response. PKR is a serine-threonine kinase comprising two conserved double-stranded RNA binding motifs (dsRBMs) in its N-terminal domain and a C-terminal kinase domain. In the cytoplasm, PKR detects dsRNA via its N-terminal dsRBM, contributing to its homodimerization and rapid autophosphorylation at residues Thr446 and Thr451 (Dauber B and Wolff T, 2009). PKR transduces signaling by phosphorylating its targeted substrate eIF2α at Ser51, abolishing the initiation of protein synthesis (Dalet A et al., 2015). Accumulated evidence has demonstrated that PKR-mediated inhibition of translation initiation plays crucial roles in human physiological and pathological processes, such as the regulation of cell proliferation, differentiation, apoptosis, viral infection, cancer and inflammation (Qiao H et al., 2021, Hsu LC et al., 2004, Feng H et al., 2017, Darini C et al., 2019, Lu B et al., 2012). However, the precise regulatory mechanism remains unclear. In addition to activating PKR, dsRNA also triggers innate immunity by binding to PRRs distributed on different organelles in cells (Liu G and Gack MU, 2020). Several sensors in the cytoplasm are known to detect dsRNA, including retinoic acid-inducible gene I (RIG-I) and melanoma differentiation associated protein 5 (MDA5) (Chen YG and Hur S, 2022). Signaling through RIG-I or MDA5 requires the mitochondrial adaptor protein MAVS (also known as IPS1, VISA, or CARDIF), which serves as an adaptor for the events of TBK1- or IKK-mediated phosphorylation of downstream cascades, including the activation of transcription factors IRF3 and NF-κB, which induces the production of IFNs and proinflammatory cytokines (Wu J and Chen ZJ, 2014, Liu S et al., 2015). The innate immune response is crucial for host defense against viral infection, while the function of the virus-induced stress response during viral infection is not clear.

TRIM21 is one of the members of the tripartite motif (TRIM) superfamily, which plays an important role in diverse biological and pathophysiological processes, including systemic autoimmune diseases, such as systemic lupus erythematosus (SLE), antiviral innate immunity and cancer proliferation (Wang L and Ning S, 2021, Lee AYS et al., 2021, Xue B et al., 2018). Our previous study showed that TRIM21 interacts with MAVS and catalyzes the K27-linked ubiquitination of MAVS to restrict viral infection by promoting the innate immune response to viral infection (Xue B et al., 2018). As one of the important PRRs, PKR is substantially activated upon viral infection. Given the important role of PKR in viral infection, whether TRIM21 regulates the PKR signaling pathway should be more comprehensively understood. In this study, we find that TRIM21 negatively regulates the PKR signaling pathway under stress, such as viral infection or thapsigargin (TG) treatment. Mechanistically, TRIM21 interacts with PP1α and promotes the K6-linked polyubiquitination of PP1α, augmenting the interaction between PKR and PP1α, which inactivates PKR and subsequently activates eIF2α, initiating protein synthesis. Moreover, proteomics analysis reveals that TRIM21 constitutively restricts viral infection by reversing PKR-mediated inhibition of the protein synthesis of intrinsic antiviral genes. The crucial regulatory role of TRIM21 in PKR activation has led to an idea that decreasing the expression of TRIM21 in cancer cell is helpful for interferon-dependent immunotherapy, and, consistent with that TRIM21 destabilizes p53 protein, TRIM21 can used as a target for tumor treatment. Collectively, our data highlight the essential role of TRIM21 in regulating protein translation, which may provide a new strategy for the treatment of translation initiation-associated diseases.

## Results

### TRIM21 is associated with PKR under viral infection

To investigate the relationship of PKR and TRIM21, we firstly analyzed the protein levels of TRIM21 and PKR in different tissues by using the database: The Human Protein Atlas (https://www.proteinatlas.org/). TRIM21 and PKR were found to be widely expressed in various tissues (Figs 1A, 1B), indicating a potential relationship between TRIM21 and PKR. Because PKR can be activated by viral infection, to further confirm the relevance of PKR and TRIM21, we analyzed the expression level of TRIM21 under various viral infections. The expression of TRIM21 was induced under various viral infections, indicating that TRIM21 may play a role under viral infection (Fig 1C). To further confirm our conclusion, we investigated the dynamic expression of TRIM21 after viral infections, including HCV, Zika virus, Dengue virus, Ebola virus, Influenza A virus, HSV-1, Human cytomegalovirus and Rrift Valley Fever virus by using multi-omics portal of viral infection (https://mvip.whu.edu.cn/) (Figs 1D, 1E). The mRNA level of TRIM21 was rapidly induced at the early time of infection, and was decreased at the later time of infection, which indicated that TRIM21 may be tightly relevant with PKR activation, because PKR is activated at the early time of viral infection. Collectively, these data indicated that TRIM21 may be associated with PKR activation in response to stress.

**Fig 1.**
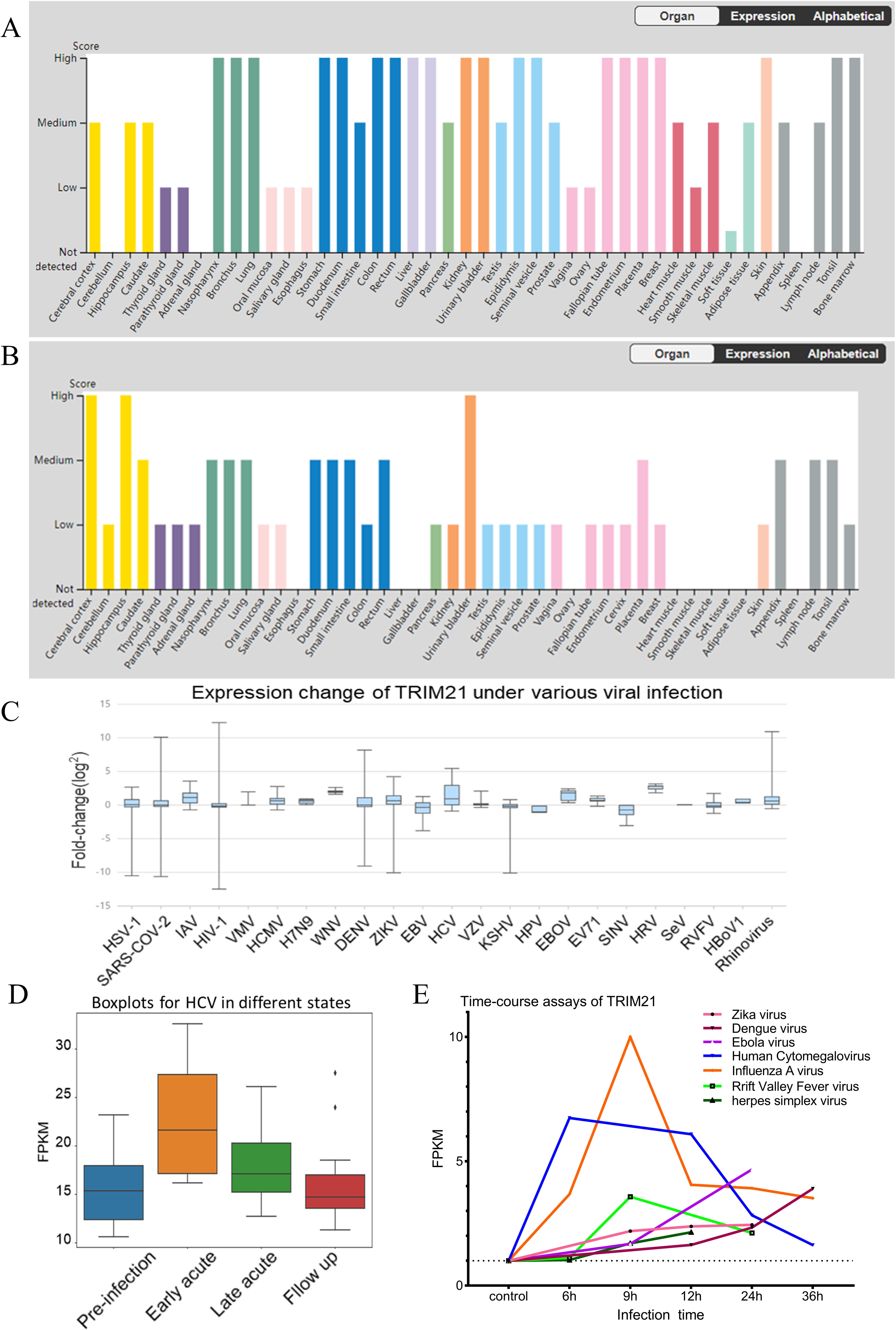
TRIM21 is associated with PKR under viral infection. A-B The dynamics of TRIM21 expression under different types of viral infections. Longitudinal transcriptomic characterization of the TRIM21 expression in different states of hepatitis C virus infection, Pre-infection baseline (Variable); Early acute (2-9 weeks, mean 6 weeks); Late acute (15 – 20 weeks, mean 17 weeks); and Follow up (25 – 71 weeks, mean 52 weeks) (A). The mRNA level of TRIM21 at different time-points after various viral infections (B). C Fold-changes of TRIM21 expression between infection and corresponding controls. D-E Protein expression levels of TRIM21and PKR in different tissues.

### TRIM21 inhibits the activation of PKR and promotes protein synthesis

To ascertain whether TRIM21 affects the PKR response under stress, we used CRISPR– Cas9 gene editing technology to establish *TRIM21*-deficient A549 cell lines (sg-*TRIM21*). TRIM21 was knocked out effectively in A549 cells (Fig EV1A). We initiated the events of PKR signaling by employing a classic agonist of PKR, poly (I:C), in these cells (Xue et al., 2018). The levels of p-PKR and p-eIF2α in *TRIM21*-deficient cells were significantly higher than those in wild-type cells responding to poly (I:C) (Fig 2A), suggesting a negative regulatory role of TRIM21 in the PKR signaling pathway. To rule out the possibility of an off-target effect of *TRIM21* sgRNA, we used shRNA specifically targeting *TRIM21* to knock down the expression of TRIM21 in A549 cells (Fig EV1B). Similarly, knockdown of *TRIM21* dramatically enhanced poly (I:C)-induced phosphorylation of PKR and eIF2α (Fig EV1C). Next, we examined the effects of TRIM21 on PKR activation upon viral infection. Vesicular stomatitis virus (VSV) or Sendia virus (SeV) infection activated PKR and eIF2α, and TRIM21 deficiency enhanced their activation (Figs 2B and 2C). Moreover, we obtained similar results in *TRIM21*-silenced cells upon VSV or SeV infection (Figs EV1D and EV1E). We performed the above experiments in HLCZ01 cells, which support the entire life cycle of hepatitis B virus (HBV) and hepatitis C virus (HCV) (Yang et al., 2014). The levels of p-PKR and p-eIF2α were increased in *TRIM21*-silenced HLCZ01 cells treated with poly (I:C) (Fig EV1F) or virus (Figs EV1G and EV1H). These data demonstrated that TRIM21 represses the activation of dsRNA- or virus-triggered PKR signaling pathways. To further investigate the negative regulatory role of TRIM21 in PKR activation, we assessed PKR activation under different stimuli. DsRNAs produced during viral infection can activate PKR (McCormick and Khaperskyy, 2017). Thus, we assessed the cellular response to interferon stimulatory DNA (ISD) stimulation or DNA virus (HSV) infection. Notably, ISD stimulation or HSV infection substantially activated PKR, while the activated PKR was inhibited by TRIM21 overexpression in the cells (Fig EV1I and Fig 2D). Moreover, the PKR activation induced by ER stress activator Thapsigargin (TG) was also repressed by TRIM21 (Sharma et al., 2017) (Figs 2E and 2F). Consistently, TRIM21 promoted global translation initiation upon viral infection or TG treatment (Fig 2G-2I). Collectively, these data supported that TRIM21 inhibits the activation of the PKR signaling pathway and the subsequent release of protein synthesis inhibition.

**Figure 2.**
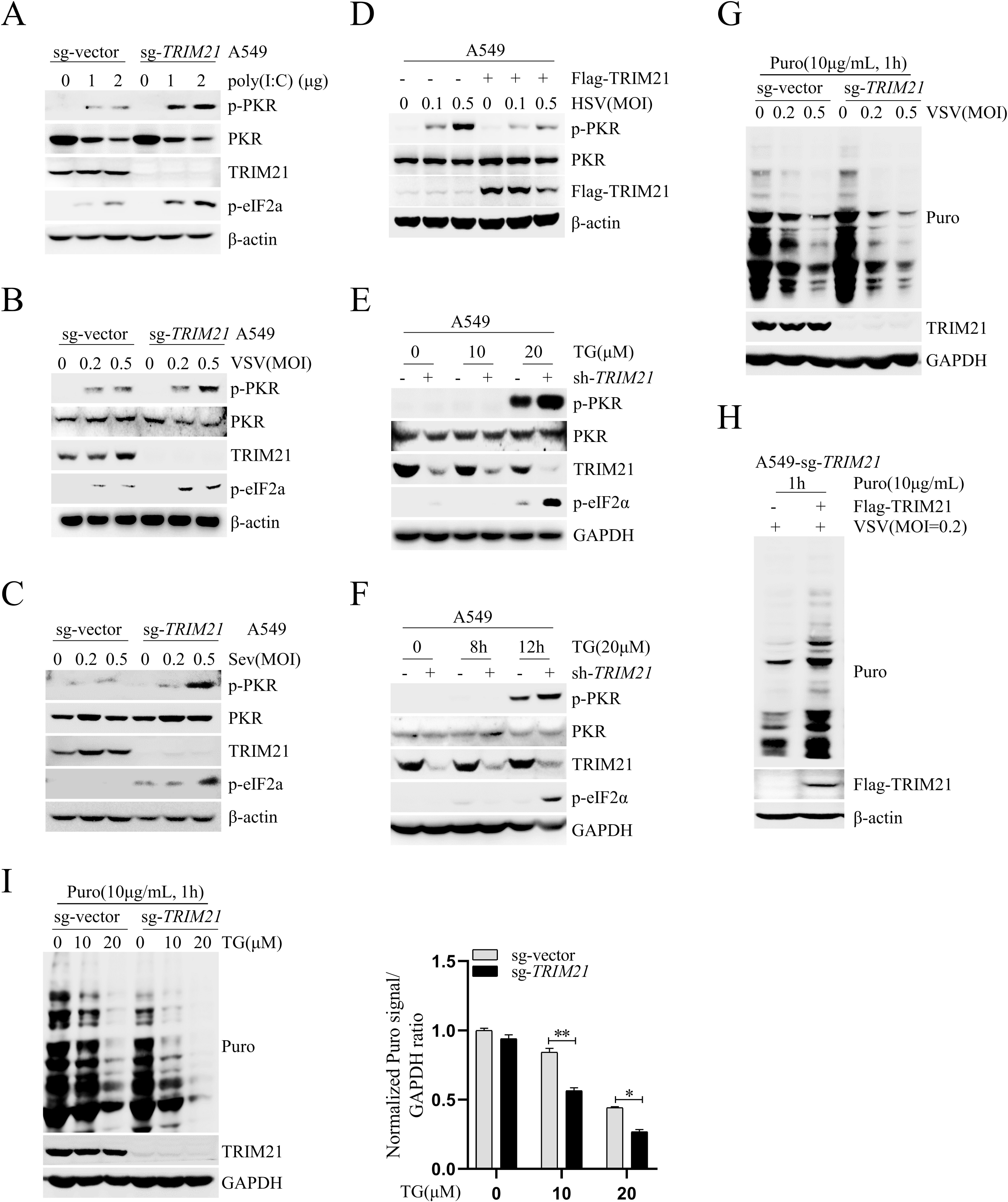
TRIM21 inhibits the activation of PKR. A-C Wild type A549 cells (sg-vector) or TRIM21-knocked out A549 cells (A549-sg-*TRIM21*) were transfected with poly(I:C) for 12 h (A), or infected with VSV (MOI=0.2 or 0.5) for 6 h (B) or Sev (MOI=0.2 or 0.5) for 12 h (C). The indicated proteins were detected by western blot. β-actin was used as an internal control. D A549 cells were transfected with p3xFlag-CMV-vector or p3xFlag-CMV-TRIM21 for 48 h, then infected with HSV (MOI=0.1 or 0.5) for 6 h. The indicated proteins were detected by western blot. β-actin was used as an internal control. E-F A549 cells were infected with lentivirus-sh-vector (sh-vector) or lentivirus-sh-TRIM21 (sh-*TRIM21*) for 48 h, then stimulated by thapsigargin (TG) (10 μM or 20 μM) for 12 h (E), or TG (20 μM) for indicated time points (F). The indicated proteins were detected by western blot. G Puromycin incorporation assays demonstrated the cellular protein synthesis in mock- or VSV-infected wild type A549 cells (sg-vector) or *TRIM21*-deficient A549 cells (sg-*TRIM21*). Cells were pulse-labeled with puromycin (10μg/mL) for 1 h prior to harvest. GAPDH was used as a control. H Puromycin incorporation assays of the cellular protein synthesis in TRIM21-deficient A549 cells (sg-*TRIM21*) transfected with p3xFlag-CMV-vector or p3xFlag-CMV-TRIM21 for 48 h, then infected by VSV (MOI=0.2) for 6 h. β-actin was used as an internal control. I Immunoblot analysis of puromycin and TRIM21 in wild type A549 cells (sg-vector) or *TRIM21*-deficient A549 cells (sg-*TRIM21*) stimulated with TG (10 μM or 20 μM) for 12 h. GAPDH was used as an internal control. Experiments were independently repeated two or three time with similar results. Student’s two-sided t test, and the data are represented as mean ± SD with three biological replicates. \**P*<0.05 versus the control; \*\**P*<0.01 versus the control.

### The ubiquitin ligase activity of TRIM21 is required for TRIM21-mediated inhibition of PKR activation

To explore the mechanism of TRIM21-mediated inhibition of the PKR signaling pathway, we performed co-IP to analyze the interaction between PKR or eIF2α and TRIM21. TRIM21 interacted with PKR but not eIF2α (Figs 3A, 3B). Furthermore, the endogenous TRIM21 and PKR interaction was confirmed in A549 cells, and their interaction was enhanced under either viral infection or TG stimulation (Figs 3C, 3D), indicating that TRIM21 may inhibit PKR activation by targeting PKR. TRIM21 is an E3 ligase with enzymatic activity in the RING finger domain. To investigate whether the E3 ligase activity of TRIM21 is involved in the regulation of PKR activation, we cotransfected pFlag-tagged PKR and hemagglutinin (HA)-tagged ubiquitin (HA-ub) together with pV5-tagged TRIM21 into HEK293T cells and found that TRIM21 was unable to catalyze the ubiquitination of PKR (Fig 3E). However, wild-type TRIM21 (TRIM21-WT), but not the E3 ligase-inactive mutant (TRIM21-C16S), in which Cys 16 was replaced with Ser 16, decreased PKR phosphorylation under VSV or TG stimulation (Figs 3F, 3G), suggesting that the E3 ligase activity of TRIM21 is required for PKR inactivation. All the data demonstrated that TRIM21 inhibits PKR activation by targeting PKR and that the E3 ligase activity of TRIM21 is crucial in this process.

**Figure 3.**
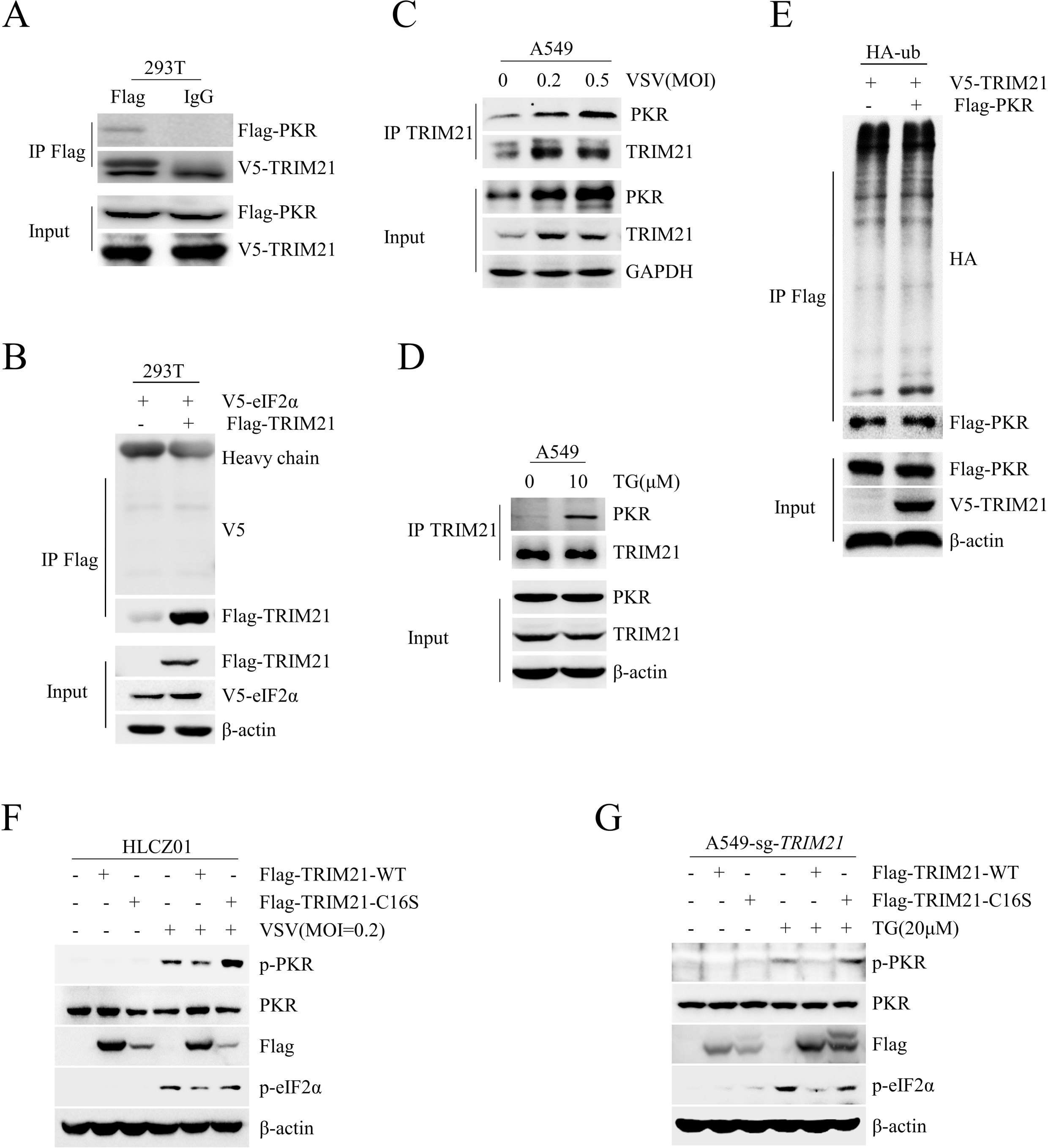
The ubiquitin ligase activity of TRIM21 is required for TRIM21-mediated inhibition of PKR activation. A p3xFlag-CMV-PKR and pcDNA3.1a-TRIM21 were co-transfected into HEK293T cells for 48 h. IP and immunoblotting assays were performed with the indicated antibodies. B HEK293T cells were co-transfected with p3xFlag-CMV-TRIM21 and pcDNA3.1a-eIF2α for 48 h. IP and immunoblotting assays were performed with the indicated antibodies. C-D Endogenous PKR and TRIM21 interaction was analyzed by IP in A549 cells with VSV (MOI=0.2 or 0.5) infection for 6 h (C) or with TG (10 μM) treatment for 12 h (D). E HEK293T cells were co-transfected with the indicated plasmids for 48 h. Ubiquitination and immunoblotting assays were performed with the indicated antibodies. F HLCZ01 cells were transfected with p3xFlag-CMV-TRIM21-WT or p3xFlag-CMV-TRIM21-C16S for 48 h, then infected with VSV (MOI=0.2) for 6 h. The indicated proteins were detected by western blot. β-actin was used as an internal control. G *TRIM21*-deficient A549 cells (sg-*TRIM21*) were transfected with p3xFlag-CMV-TRIM21-WT or p3xFlag-CMV-TRIM21-C16S for 48 h, then stimulated with TG (20 μM) for 12 h. The indicated proteins were detected by western blot. β-actin was used as an internal control. Experiments were independently repeated two or three time with similar results.

### TRIM21 represses PKR activation by targeting PP1α

PKR is a serine–threonine kinase that comprises a kinase domain and two dsRNA binding domains regulating its activity. Upon engagement with dsRNA in the cytoplasm, PKR undergoes homodimerization and subsequent rapid autophosphorylation at Thr446 and Thr451, increasing its catalytic activity (Dauber and Wolff, 2009). Therefore, there are three steps in PKR activation: dsRNA sensing, homodimerization and autophosphorylation. To determine which step of PKR activation is regulated by TRIM21, we performed RIP analysis to explore whether TRIM21 affects PKR binding to viral RNA in HEK293T cells infected with VSV. PKR constitutively bound to viral RNA, while TRIM21 knockdown did not abolish their interaction (Fig 4A), suggesting that TRIM21 has no effect on the detection of dsRNA by PKR. To test whether TRIM21 affects PKR homodimerization, we delivered pFlag-tagged PKR and pMyc-tagged PKR together with different doses of pV5-tagged TRIM21 into HEK293T cells. TRIM21 could not abolish the interaction between Flag-tagged PKR and pMyc-tagged PKR (Fig 4B), indicating that TRIM21 may not suppress homodimerization. These data indicated that TRIM21 may regulate PKR activation by impairing its phosphorylation. Given that PKR phosphorylates itself, we speculated that TRIM21 may target the phosphatases of PKR to inactive PKR. PP1α is the key phosphatase of PKR (Tan et al., 2002). To investigate whether TRIM21 targets PP1α to regulate PKR, we performed co-IP assays to analyze the interaction between PP1α and TRIM21 as well as that of PP1α with PKR in HEK293T cells. Indeed, PP1α interacted with both TRIM21 and PKR (Fig 4C). The endogenous PP1α and TRIM21 interaction was confirmed in A549 cells, and stress, including VSV infection or TG stimulation, augmented their interaction (Figs 4D, 4E). Moreover, an in vitro pull-down assay with purified recombinant proteins demonstrated a direct interaction between TRIM21 and PP1α (Fig 4F). To determine which domain of TRIM21 is the key domain for PP1α binding, we constructed a series of TRIM21 truncations and cotransfected the individual truncations with PP1α into HEK293T cells. Co-IP assay results showed that the PRY-SPRY domain of TRIM21 was essential for its interaction with PP1α (Figs 4G, 4H). All the data supported that TRIM21 inhibits PKR activation by targeting PP1α.

**Figure 4.**
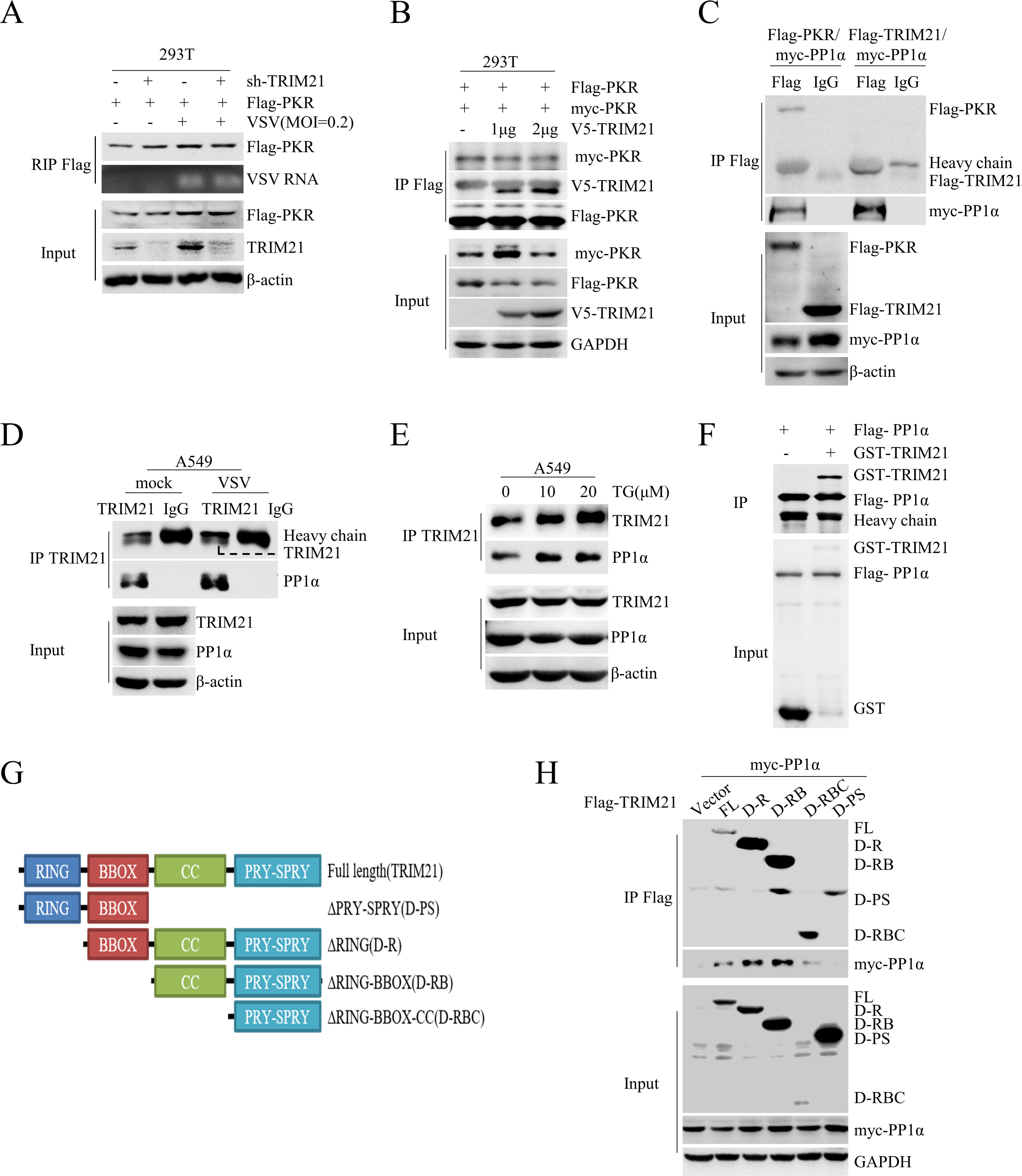
TRIM21 represses PKR activation by targeting on PP1α. A HEK293T cells preinfected with lentivirus-sh-vector (sh-vector) or lentivirus-sh-*TRIM21* (sh-*TRIM21*) for 24 h were transfected with p3xFlag-CMV-vector or p3xFlag-CMV-PKR for 48 h, then infected VSV (MOI=0.2) for 6 h. RIP assay was performed to test the VSV RNA binding to PKR. B p3xFlag-CMV-PKR, pCMV-N-Myc-PKR and pcDNA3.1a-TRIM21 were co-transfected into HEK293T cells for 48 h. IP and immunoblotting assays were performed with the indicated antibodies. C IP analysis of the interaction of PP1α and TRIM21 as well as PP1α and PKR in HEK293T cells co-transfected with the indicated plasmids for 48 h. D-E Endogenous PP1α and TRIM21 interaction was analyzed by IP in A549 cells infected with VSV (MOI=0.2) for 6 h (D) or stimulated with TG (10 μM or 20 μM) for 12 h (E). F IP analysis of the interaction between GST-TRIM21 protein and Flag- PP1α protein purified from bacteria. G Schematic illustration of TRIM21 truncations. H IP analysis of the interaction between individual domain of TRIM21 and PP1α in HEK293T cells co-transfected with p3xFlag-CMV-TRIM21 truncations and pCMV-N-Myc-PP1α for 48 h. Experiments were independently repeated two or three time with similar results.

### TRIM21 promotes PP1α polyubiquitination

Since TRIM21 targets PP1α and the regulation of PKR by TRIM21 depends on its ubiquitin ligase activity, we investigated whether the E3 ligase activity of TRIM21 targets PP1α. We delivered pTRIM21-WT or pTRIM21-C16S together with pFlag-tagged PP1α and pHA-ub into HEK293T cells and found increased ubiquitination of PP1α in TRIM21-WT-transfected cells but not in TRIM21-C16S-transfected cells (Fig 5A), suggesting that TRIM21 promotes the ubiquitination of PP1α. Different types of polyubiquitin linkages have distinct functions. We cotransfected pPP1α into HEK293T cells with individual ubiquitin mutants (K6O, K11O, K27O, K29O, K33O, K48O or K63O), each of which contained only one lysine residue available for modification. TRIM21 specifically promoted K6-linked ubiquitination of PP1α (Fig 5B). Moreover, TRIM21-WT but not TRIM21-C16S catalyzed the K6-linked ubiquitination of PP1α. As a negative control, neither TRIM21-WT nor TRIM21-C16S promoted K11-linked ubiquitination of PP1α (Fig 5C), which further confirmed that the K6-linked ubiquitination of PP1α was mediated by TRIM21. Next, we examined which lysine residue of PP1α is ubiquitinated by TRIM21. PP1α was divided into two fragments, the N-terminus and C-terminus, and either the N-terminus or C-terminus of PP1α was transfected together with TRIM21-WT or TRIM21-C16S as well as HA-ub into HEK293T cells (Fig 5D). Co-IP assays revealed that the N-terminus of PP1α was ubiquitinated by TRIM21 (Fig 5E). Since the N-terminus of PP1α contains 5 lysine residues, we generated the mutant PP1α-K0, in which all of the lysine residues in the N-terminus of PP1α were replaced with arginine. Then, we reintroduced individual lysine residues into PP1α-K0 to generate single lysine mutants (Fig 5F). Co-IP assay results showed that the K60 mutant of PP1α was sufficient for TRIM21-mediated polyubiquitination of PP1α (Fig 5G), suggesting that TRIM21 may promote the K6-linked ubiquitination of PP1α on Lys60. Furthermore, TRIM21 lost the ability to catalyze K6-linked ubiquitination of the K60R mutant of PP1α, in which lysine 60 was replaced by arginine 60 (Fig 5H). Collectively, these data demonstrated that TRIM21 promotes the K6-linked ubiquitination of PP1α at Lys60.

**Figure 5.**
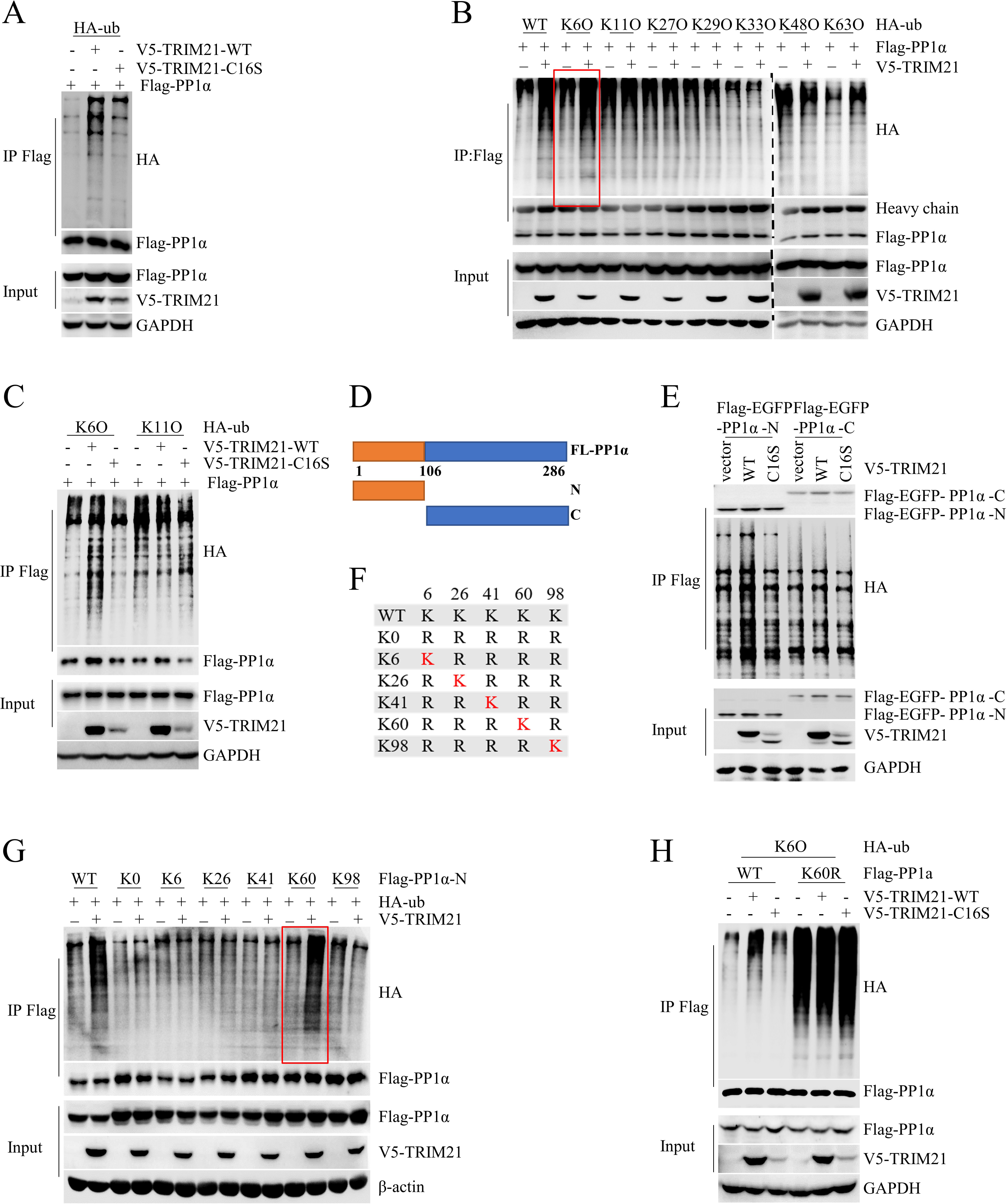
TRIM21 promotes the polyubiquitination of PP1α. A IP analysis of the ubiquitination of PP1α in HEK293T cells co-transfected with the indicated plasmids for 48 h. B HEK293T cells were co-transfected with p3xFlag-CMV-PP1α and pcDNA3.1a-TRIM21 as well as the indicated HA-tagged ubiquitin mutants for 48 h. Ubiquitination and immunoblotting assays were performed with the indicated antibodies. C HEK293T cells were co-transfected with the indicated plasmids for 48 h. Ubiquitination and immunoblotting assays were performed with the indicated antibodies. D A schematic diagram of PP1α truncations. E IP analysis of the ubiquitination of C terminus and N terminus of PP1α in HEK293T cells co-transfected the indicated plasmids for 48h. F-G TRIM21 promotes the polyubiquitination of PP1α on Lys60. Mutants of only one lysine residue retained within N-terminal PP1α (F). HEK293T cells were co-transfected with pcDNA3.1a-vector, pcDNA3.1a-TRIM21 or the indicated mutants of p3xFlag-CMV-PP1α-N for 48 h. Ubiquitination and immunoblotting were performed with the indicated antibodies (G). H TRIM21 promotes K6-linked ubiquitination of PP1α on Lys60. HEK293T cells were co-transfected with the indicated plasmids for 48 h. Ubiquitination and immunoblotting assays were performed with the indicated antibodies. Experiments were independently repeated two or three time with similar results.

### TRIM21 inhibits the activation of the PKR signaling pathway by promoting the K6-linked ubiquitination of PP1α

Since our data showed that the E3 ligase activity of TRIM21 is sufficient for PKR inactivation and that TRIM21 catalyzes the K6-linked ubiquitination of PP1α, we speculated that the inhibition of PKR activation by TRIM21 may depend on PP1α. To determine the role of PP1α in the regulation of PKR activation by TRIM21, we knocked down the expression of *TRIM21* and *PP1α* in A549 cells. TRIM21 knockdown substantially enhanced PKR activation upon VSV infection, while the phenomenon was lost in *PP1α*-silenced cells (Fig 6A), indicating that TRIM21 may inhibit PKR activation via PP1α. Next, we constructed a mutant with inactive PP1α phosphatase activity (PP1α-H248K). TRIM21 lost the ability to inactivate PKR by PP1α-H248K introduction upon VSV infection or TG stimulation (Fig 6B-6D). Next, we explored whether K6-linked ubiquitination of PP1α is essential for TRIM21-mediated PKR inactivation. We reintroduced the wild-type PP1α (PP1α-WT) or the K60 mutant PP1α (PP1α-K60R) together with Flag-tagged TRIM21 into PP1α-silenced cells to detect the phosphorylation of PKR. TRIM21 inhibited PKR phosphorylation in PP1α-WT-transfected cells but not in PP1α-K60R-transfected cells under VSV or TG stimulation (Figs 6E, 6F), indicating that TRIM21-mediated ubiquitination of PP1α is crucial for PKR inactivation. Collectively, these data suggest that TRIM21 impairs PKR activation by promoting the K6-linked ubiquitination of PP1α.

**Figure 6.**
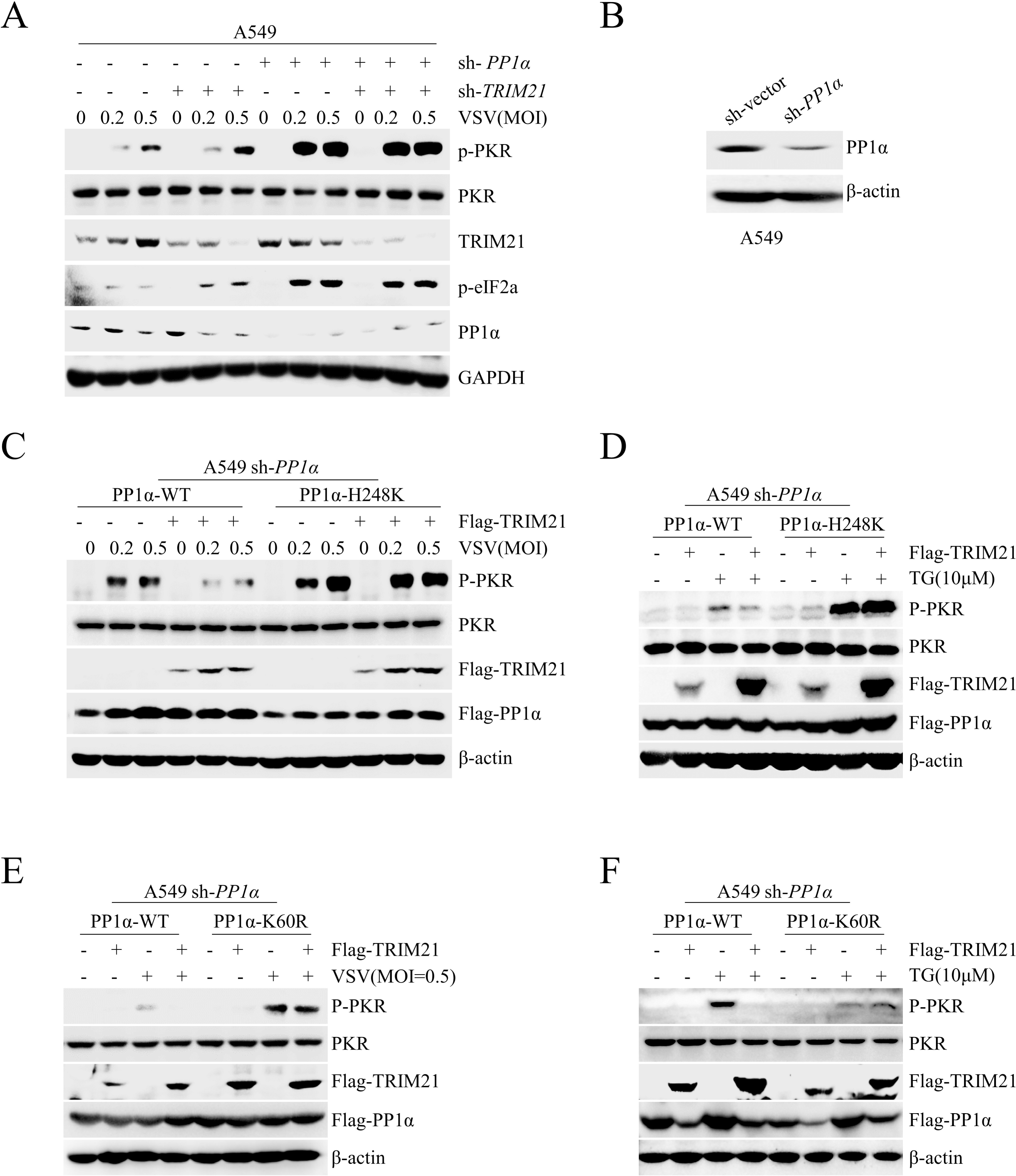
TRIM21 inhibits PKR signaling pathway activation via PP1α. A *PP1α-*silenced A549 cells were infected with lentivirus-sh-vector (sh-vector) or lentivirus-sh-*TRIM21* (sh-*TRIM21*) for 48 h, then infected by VSV (MOI=0.2 or 0.5) for 6 h, The activation of PKR was analyzed by western blot. GAPDH was used as an internal control. B Immunoblot analysis detected PP1α protein in wild type A549 cells (sh-vector) or *PP1α*-silenced A549 cells (sh-*PP1α*). β-actin was detected as an internal control. C-D *PP1α*-silenced A549 cells (sh-*PP1α*) pre-transfected with p3xFlag-CMV-PP1α-WT or p3xFlag-CMV-PP1α-H248K for 24 h were infected with lentivirus-TRIM21 for 48h, then infected with VSV (MOI=0.2 or 0.5) for 6 h (C) or stimulated with TG (10 μM) for 12 h (D). E-F *PP1α*-silenced A549 cells (sh-*PP1α*) were transfected with p3xFlag-CMV-PP1α-WT or p3xFlag-CMV-PP1α-K60R for 48 h, then infected with VSV (MOI=0.5) for 6 h E or stimulated with TG (10 μM) for 12 h (F). The activation of PKR was analyzed by western blot. β-actin was detected as an internal control. Experiments were independently repeated two or three time with similar results.

### TRIM21 inhibits PKR activation by enhancing the PKR-PP1α interaction

To determine how TRIM21-mediated polyubiquitination of PP1α regulates the activation of PKR, we examined whether TRIM21 affects the stability of the PP1α protein. TRIM21 did not alter the level of PP1α protein under VSV infection or TG treatment (Fig 7A-7C), suggesting that TRIM21 does not affect the stability of PP1α protein. PP1α interacts with PKR and subsequently dephosphorylates phosphorylated PKR, so we performed a co-IP assay to examine whether TRIM21 affects the interaction between PKR and PP1α under different stimuli. Although TRIM21 had no effect on the interaction of PKR and PP1α in the resting state, *TRIM21* knockdown significantly reduced their interaction upon VSV infection or TG stimulation (Figs 7D, 7E). These data indicated that TRIM21 inhibits PKR activation by enhancing the interaction between PKR and PP1α under stress. Moreover, stress augmented the interaction between PKR and PP1α in TRIM21-WT-reconstituted cells but not in TRIM21-C16S-reconstituted cells upon VSV infection (Fig 7F), suggesting that TRIM21-mediated ubiquitination of PP1α is essential for the enhancement of the PKR-PP1α interaction. In summary, extracellular stimuli, such as viral infection or TG treatment, promotes the interaction between TRIM21 and PP1α and subsequent PP1α ubiquitination by TRIM21. Ubiquitination of PP1α enhances its interaction with PKR, which promotes dephosphorylation of PKR and repression of PKR and subsequent eIF2α activation, thereby initiating protein translation (Fig 7G).

**Figure 7.**
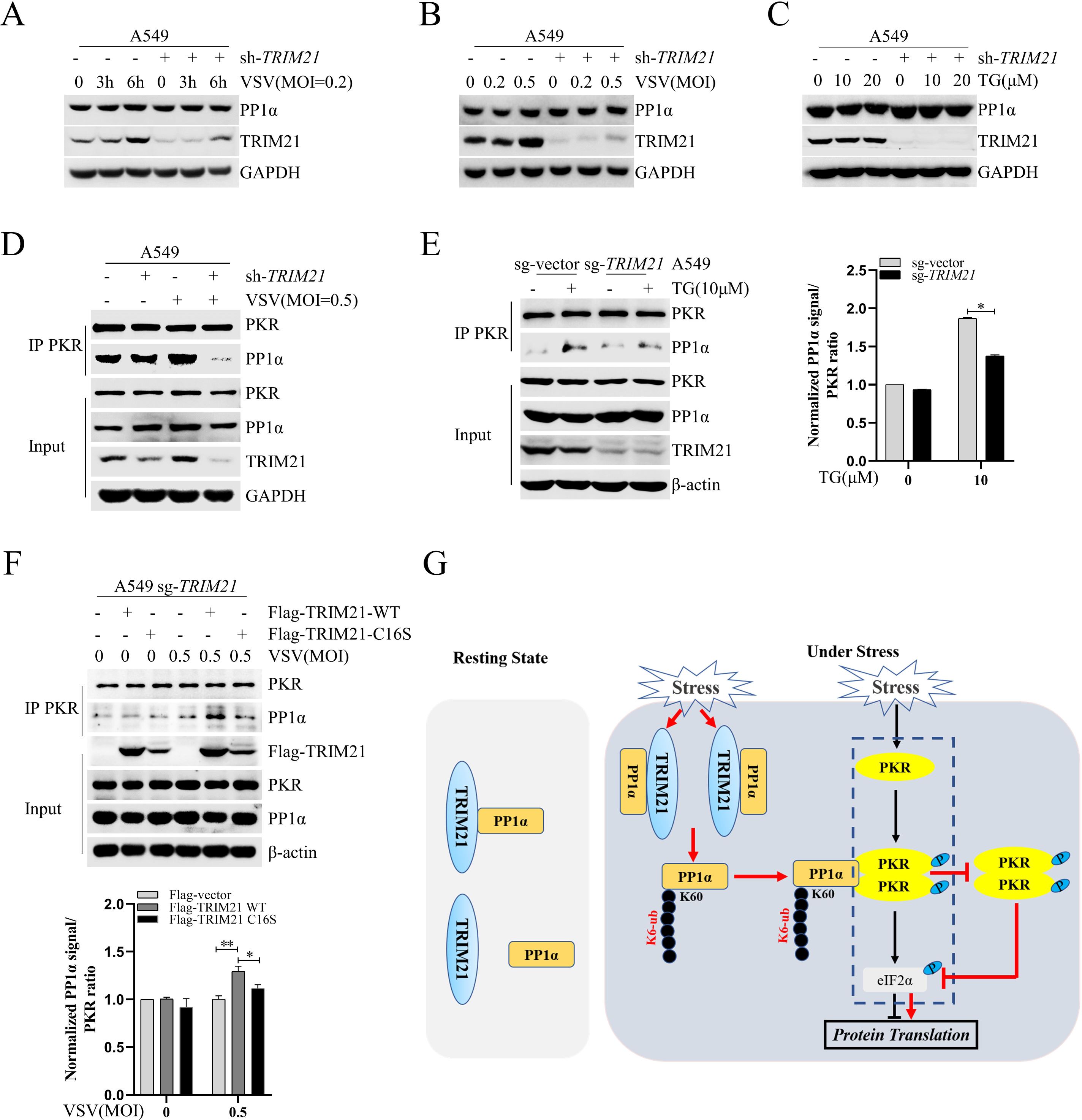
TRIM21 inhibits PKR activation by augmenting PKR-PP1α interaction. A-B A549 cells were infected with lentivirus-sh-vector (sh-vector) or lentivirus-sh-*TRIM21* (sh-*TRIM21*) for 48 h, then infected by VSV (MOI=0.2) for indicated times (A), or VSV (MOI=0.2 or 0.5) for 6 h (B). PP1α protein was analyzed by western blot. GAPDH was used as an internal control. C A549 cells were infected with lentivirus-sh-vector (sh-vector) or lentivirus-sh-*TRIM21* (sh-*TRIM21*) for 48 h, then stimulated with TG (10μM or 20μM) for 12 h. PP1α protein was analyzed by western blot. GAPDH was used as an internal control. D-E TRIM21 blocks PKR-PP1α interaction. A549 cells were infected with lentivirus-sh-vector (sh-vector) or lentivirus-sh-*TRIM21* (sh-*TRIM21*) for 48 h, then infected by VSV (MOI=0.5) for 6 h. The interaction of PP1α and PKR was analyzed by IP and western blot (D). The wild type A549 cells (sg-vector) or *TRIM21*-deficient A549 cells (sg-*TRIM21*) were stimulated with TG (10 μM) for 12 h, The interaction of PP1α and PKR was analyzed by IP and western blot (E). F *TRIM21*-deficient A549 cells (sg-*TRIM21*) were transfected with p3xFlag-CMV-TRIM21-WT or p3xFlag-CMV-TRIM21-C16S, then infected by VSV (MOI=0.5) for 6 h. The interaction of PP1α and PKR was analyzed by IP and western blot. G Schematic model of TRIM21 regulating the translation initiation. Upon stress stimulation, TRIM21 augments the polyubiquitination of PP1α, enhancing the PKR-PP1α interaction, leading to PKR inactivation and subsequent release of protein synthesis inhibition. Experiments were independently repeated two or three time with similar results. Student’s two-sided t test, and the data are represented as mean ± SD with three biological replicates. \**P*<0.05 versus the control.

### TRIM21 restricts viral infection by inhibiting PKR activation

Several studies have demonstrated that PKR activation is essential for viral escape (Feng et al., 2017, Garaigorta and Chisari, 2009). We speculated that TRIM21 has the ability to restrict viral infection by inhibiting PKR-mediated translational shutdown. Because TRIM21 is able to defend against viral infection by promoting IFN production, we knocked out *IFNAR1* in both A549 cells (A549-sg-*IFNAR1*) and HLCZ01 cells (HLCZ01-sg-*IFNAR1*) (Fig EV2A). Notably, the levels of phosphorylated PKR and eIF2α were repressed by TRIM21 in the cells upon VSV infection (Figs 8A, 8B). Consistently, silencing TRIM21 significantly increased the replication and production of the virus in the cells (Figs 8C, 8D). The antiviral role of TRIM21 was further confirmed in A549-sg-*IFNAR1* cells infected with SeV (Fig EV2B). These data indicated that TRIM21 can restrict viral infection in an IFN-independent manner. To exclude the possible role of type-III IFN-mediated antiviral response by viral infection, we knocked down STAT2 to abolish the type-III IFN signaling pathway in A549-sg-*IFNAR1* cells (Fig EV2C). The replication of VSV was increased in *TRIM21*-silenced cells (Fig 8E), indicating that the restriction of viral infection by TRIM21 does not rely on type III IFNs. Furthermore, we performed the investigation by abolishing IFN production. Silencing *TRIM21* augmented viral replication and PKR activation in *IRF3*-knockdown cells upon VSV infection (Figs EV2D, 8F and 8G). Similarly, in Huh7.5 cells, in which one of the main receptors for RNA viruses, RIG-I, is deficient, the replication of VSV and SeV and the phosphorylation of PKR and eIF2α were increased in *TRIM21*-silenced cells (Figs 8H, 8I and Fig EV2E). These data demonstrated that TRIM21 restricts viral infection independent of its promotion of innate immunity. The antiviral function of TRIM21 was further confirmed in mice. We knocked down *Trim21* in mice by intravenous injection of the *Trim21*-shRNA lentivirus. Indeed, knockdown of *Trim21* significantly augmented the replication of VSV in the liver, spleen and lung tissues in the mice (Figs 8J, 8K). Next, we examined whether TRIM21 restricts viral infection by inhibiting PKR activation. *IFNAR1-PKR* double-deficient A549 cell lines were constructed (Fig EV2F). Depleting *TRIM21* increased VSV replication in the cells. However, the antiviral function of TRIM21 was lost in *IFNAR1-PKR* double-deficient cells (Fig 8L). Likewise, PKR-deficient Huh7.5 cells with knockdown of *TRIM21* showed similar VSV replication to that of the control cells (Figs EV2G, 2M). We obtained similar results when cells were subjected to SeV infection (Fig EV2H). Moreover, Extended Data expression of TRIM21 in *PKR-TRIM21* double-deficient Huh7.5 cells had no effect on viral replication (Figs EV2I, EV2J). These data suggested that the antiviral function of TRIM21 relies on PKR.

**Figure 8.**
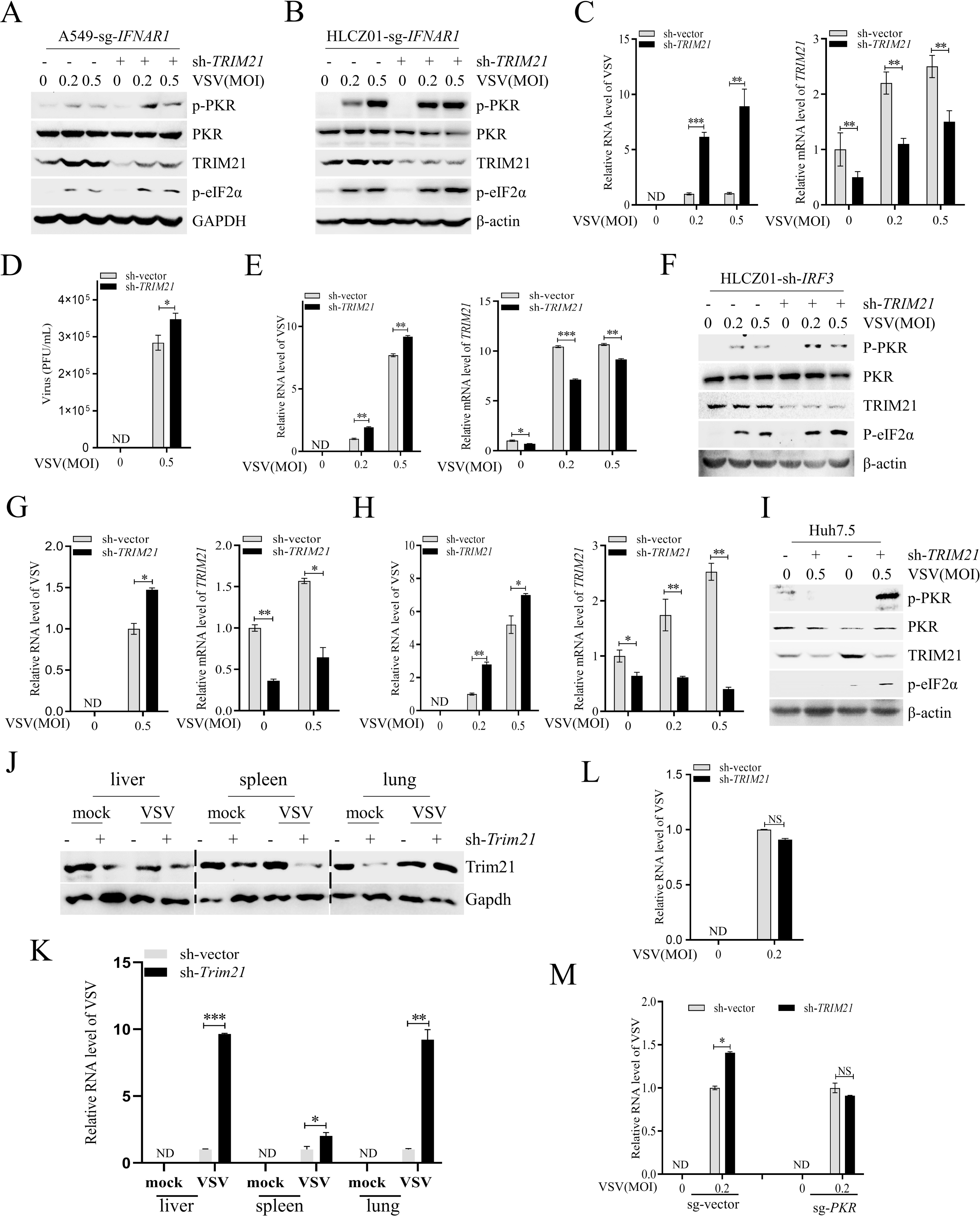
TRIM21 restricts viral infection by inhibiting the activation of PKR signaling pathway. A-B *IFNAR1*-deficient A549 cells (A) or HLCZ01 (B) cells were infected with lentivirus-sh-vector (sh-vector) or lentivirus-sh-*TRIM21* (sh-*TRIM21*) for 48 h, then infected with VSV (MOI=0.2 or 0.5) for 6 h. The indicated proteins were detected by western blot. C RT-qPCR analysis of the VSV RNA or the mRNA levels of *TRIM21* in *IFNAR1*-deficient A549 cells (sg-*IFNAR1*) infected with lentivirus-sh-vector (sh-vector) or lentivirus-sh-*TRIM21* (sh-*TRIM21*) for 48 h, then infected with VSV (MOI=0.2 or 0.5) for 6 h. D Plaque assay analysis of the VSV titer in *IFNAR1*-deficient HLCZ01 cells infected with lentivirus-sh-vector (sh-vector) or lentivirus-sh-*TRIM21* (sh-*TRIM21*) for 48 h, then infected with VSV(MOI=0.5) for 6 h. E *IFNAR1-STAT2* doubly deficient A549 cells were infected with lentivirus-sh-vector (sh-vector) or lentivirus-sh-*TRIM21* (sh-*TRIM21*) for 48 h, then infected with VSV (MOI=0.2 or 0.5) for 6 h. RT-qPCR analysis of the VSV RNA or the mRNA level of *TRIM21*. F HLCZ01-sh-*IRF3* cells were infected with lentivirus-sh-vector (sh-vector) or lentivirus-sh-*TRIM21* (sh-*TRIM21*) for 48 h, then infected with VSV (MOI=0.2 or 0.5) for 6 h. The indicated proteins were detected by western blot. G RT-qPCR analysis of the VSV RNA or the mRNA level of *TRIM21* in *IRF3*-silenced HLCZ01 cells (HLCZ01-sh-*IRF3*) infected with lentivirus-sh-vector (sh-vector) or lentivirus-sh-*TRIM21* (sh-*TRIM21*) for 48 h, then infected with VSV (MOI=0.5) for 6 h. H-I RT-qPCR analysis of the level of VSV RNA or *TRIM21* mRNA (H) or western blot analysis the indicated proteins (I) in Huh7.5 cells infected with lentivirus-sh-vector (sh-vector) or lentivirus-sh-*TRIM21* (sh-*TRIM21*) for 48 h, then infected with VSV (MOI=0.2 or 0.5) for 6 h. J Immunoblot analysis of the protein level of Trim21 in the liver, spleen and lung tissues from the eight-week-old C57BL/6 mice intravenous injected with lentivirus-sh-vector or lentivirus-sh-*Trim21* (1x10^11^ PFU/g body weight). K RT-qPCR analysis of VSV RNA in the liver, spleen and lung tissues from lentivirus-sh-vector or lentivirus-sh-*Trim21* injected eight-week-old C57BL/6 mice challenged with VSV (1x10^8^ pfu/g body weight) for 36 h. L *IFNAR1-PKR* doubly deficient A549 cells were infected with lentivirus-sh-vector (sh-vector) or lentivirus-sh-*TRIM21* (sh-*TRIM21*) for 48 h, then infected with VSV (MOI=0.2) for 6 h. The levels of VSV RNA were examined by RT-qPCR. M *PKR*-deficient Huh7.5 cells were infected with lentivirus-sh-vector (sh-vector) or lentivirus-sh-*TRIM21* (sh-*TRIM21*) for 48 h, then infected with VSV (MOI=0.2) for 6 h. The levels of VSV RNA were examined by RT-qPCR. Experiments were independently repeated two or three time with similar results. Student’s two-sided t test, and the data are represented as mean ± SD with three biological replicates. \**P*<0.05, \*\**P*<0.01, \*\*\**P*<0.001 versus the control.

### TRIM21 restricts viral infection by releasing PKR-mediated inhibition of the protein synthesis of intrinsic antiviral genes

Previous studies reported that PKR-mediated inhibition of the protein synthesis of effectors with antiviral efficiency contributes to viral escape (Feng et al., 2017, Garaigorta and Chisari, 2009). We speculated that the constitutive antiviral activities of TRIM21 may depend on its function in the impairment of PKR-mediated protein synthesis inhibition of antiviral effectors. To address our speculation, we first assessed whether TRIM21 is beneficial for the inhibition of HCV replication in response to IFN because it has been reported that PKR activation-mediated inhibition of antiviral ISG protein synthesis contributes to HCV resistance to IFN treatment (Garaigorta and Chisari, 2009). We performed the investigation in Huh7.5 cells infected by JFH-1, HCV genotype 2a, in response to IFN treatment. Indeed, TRIM21 deficiency augmented IFN-mediated PKR activation and subsequently inhibited the protein synthesis of ISGs, such as ISG15 and STAT2, in HCV-infected Huh7.5 cells (Fig EV3A). Consistently, levels of HCV RNA, NS3 protein and viral particles in the supernatant were upregulated in TRIM21-silenced cells in response to IFN-α (Fig EV3B- EV3D). Similarly, TRIM21 inhibited the replication of H77-S, HCV genotype 1a, in response to IFN (Fig EV3E). These data indicated that TRIM21 restricts HCV infection by reversing PKR-mediated inhibition of the protein synthesis of ISGs.

To further investigate whether the constitutive antiviral role of TRIM21 is dependent on its reversal of PKR-mediated translational inhibition of antiviral effectors, we performed proteomic analysis to identify the specific TRIM21-PKR-regulated antiviral effectors in *IFNAR1*-deficient cells as well as *IFNAR1/TRIM21* double-deficient cells upon VSV infection (Fig 9A). We first investigated whether TRIM21 affects global protein synthesis in *IFNAR1*-deficient cells by detecting puro-labeled nascent peptides. Consistent with the results above that TRIM21 inhibits PKR signaling pathway activation, VSV-induced nascent protein synthesis was increased by Extended Data expression of TRIM21 in *IFNAR1-TRIM21* double-deficient cells (Figs EV3F-EV3H), indicating that TRIM21 promotes protein synthesis in *IFNAR1*-deficient cells. Next, the proteomics assay showed that the protein abundances of 53 genes with significant differences were downregulated in *IFNAR1*-deficient cells with *TRIM21* depletion relative to those of *IFNAR1*-deficient cells (Fig 9A). Notably, these genes included known antiviral effectors, such as *PSMB9*, *NFKB2, OTUD4, MECP2, H2AX, NCOA7, SOD-2, S100A2*, *RFTN1, S100A4*, *FGF2, TLL2*, and *TNFAIP2* (marked with yellow) (Yamane et al., 2019, Lu et al., 2021, Liuyu et al., 2019, Cronk et al., 2017, Jha et al., 2014, Doyle et al., 2018, Wang et al., 2017, Koga et al., 2018, Tatematsu et al., 2013, Yang et al., 2020, Wang et al., 2018, Galanina et al., 2018, Chevrier et al., 2011), in addition to multiple genes with no previously recognized antiviral function (marked with red and green). Among the genes with unknown antiviral function, we focused on nine genes (marked with green) that were significantly downregulated in *TRIM21* knockout cells. A549 cells were infected with lentivirus expressing shRNAs targeting IgG Fc-binding protein (*FCGBP*), interleukin-1 receptor-associated kinase-like 2 (*IRAK2*), proteasome subunit beta type-9 (*PSMB9*), reactive oxygen species modulator 1 (*ROMO1*), latexin (*LXN*), Jade family PHD finger 1 (*JADE1*), ribonucleotide reductase regulatory TP53 inducible subunit M2B (*RRM2B*), non-SMC condensin II complex subunit H2 (*NCAPH2*), or capping actin protein (*CAPG*) and then infected with VSV. Knockdown of these genes, but not of *CAPG,* enhanced the replication of VSV in the cells (Fig 9B). We observed similar results in VSV-infected Huh7 cells and even apparent antiviral activity of *CAPG* in the cells (Fig 9C). Of note, these genes constitutively restrict HCV replication in HCV-infected Huh7.5 cells (Fig 9D). However, *FCGBP* had no effect on the replication of VSV or HCV in hepatocytes, which demonstrated that the function of *FCGBP* in hepatocytes is distinctive. These data supported that reversing PKR-mediated inhibition of the protein synthesis of antiviral genes by TRIM21 is sufficient for viral clearance.

**Figure 9.**
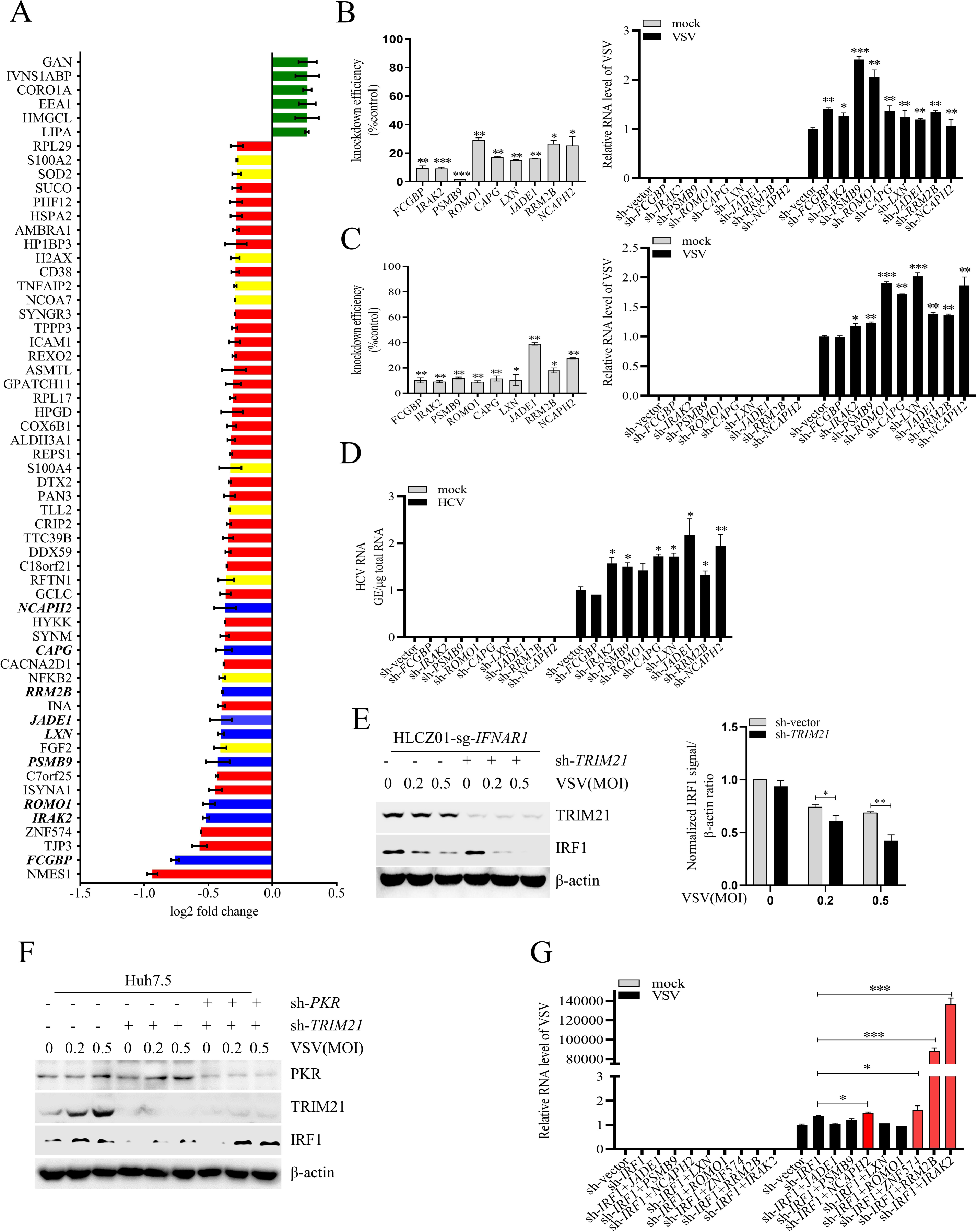
TRIM21 resists viral infection through inhibiting PKR activation-mediated global translation shutdown. A Proteome analysis of the reduced genes in *IFNAR1*-deficient A549 cells (sg-*IFNAR1*) by knockdown of *TRIM21*. The cells were infected with VSV(MOI=0.2) for 6 h. Known antiviral effectors were marked with yellow and the effectors with antiviral function verified in our study were marked with blue. B-C RT-qPCR analysis of VSV RNA or the mRNA levels of the indicated genes in A549 cells (B) or Huh7 cells (C) infected with lentivirus expressing shRNAs targeting the indicated genes for 48 h, then infected with VSV (MOI=0.2) for 6 h. D Huh7.5 cells were infected with lentivirus expressing shRNAs targeting the indicated genes for 12 h, then infected with HCV (MOI=0.1) for 72 h. HCV RNA level was analyzed by RT-qPCR. E *IFNAR1*-deficient HLCZ01 cells (HLCZ01-sg-*IFNAR1*) were infected with lentivirus-sh-vector (sh-vector) or lentivirus-sh-*TRIM21* (sh-*TRIM21*) for 48 h, then infected with VSV (MOI=0.2 or 0.5) for 6 h. IRF1 protein was analyzed by western blot. β-actin was detected as control. F Huh7.5 cells were infected with lentivirus-sh-vector (sh-vector) or lentivirus-sh-*PKR* (sh-*PKR*), and lentivirus-sh-vector (sh-vector) or lentivirus-sh-*TRIM21* (sh-*TRIM21*) for 48 h, then infected with VSV (MOI=0.2 or 0.5) for 6 h. IRF1 protein was analyzed by western blot. β-actin was detected as control. G Huh7.5 cells were infected with lentivirus expressing shRNAs targeting the indicated genes for 48 h, then infected with VSV (MOI=0.2) for 6 h. The levels of VSV RNA were examined by RT-qPCR. Experiments were independently repeated two or three time with similar results. Student’s two-sided t test, and the data are represented as mean ± SD with three biological replicates. \**P*<0.05 versus the control; \*\**P*<0.01 versus the control.

### TRIM21 restricts viral infection through IRF1-dependent and IRF1-independent mechanisms

It has been reported that basal expression of interferon regulatory factor 1 (IRF1) drives hepatocyte resistance to multiple RNA viruses by maintaining constitutive transcription of antiviral effectors, while the abundance of IRF1 protein is tightly regulated by PKR-mediated shutdown of global translation (Feng et al., 2017, Yamane et al., 2019, Wang et al., 2020). Given that TRIM21 suppresses PKR activation under stress, we hypothesized that TRIM21 can resist viral infection by promoting the protein synthesis of IRF1. Notably, silencing *TRIM21* reduced IRF1 protein levels in *IFNAR1*-deficient HLCZ01 cells infected with VSV (Fig 9E), suggesting that TRIM21 can increase the abundance of IRF1 protein. Similarly, the levels of IRF1 protein were reduced in *TRIM21*-silenced Huh7.5 cells with VSV infection, and IRF1 protein levels were restored by additional knockdown of PKR (Fig 9F), indicating that TRIM21 promotes the protein synthesis of IRF1 by negatively regulating PKR activation. Next, we examined whether the antiviral role of TRIM21 is dependent on IRF1 by depleting IRF1 in Huh7 cells. Compared with the results of single knockdown of the upregulated genes (Fig 9C), several genes, such as *JADE1*, *PSMB9*, *LXN* and *ROMO*, lost their antiviral ability after IRF1 ablation (Fig 9G), indicating an IRF1-dependent antiviral role of TRIM21. However, IRF1 had no effect on the antiviral function of *NCAPH2*, *ZNF5574*, *RRM2B* and *IRAK2* (marked with red), indicating an IRF1-independent antiviral role of TRIM21 (Fig 9G). Some genes are regulated by both TRIM21 and IRF1, contributing to the IRF1-dependent antiviral response. However, other genes, such as *NCAPH2*, *ZNF5574*, *RRM2B* and *IRAK2*, were regulated by TRIM21 but not IRF1, causing an IRF1-independent antiviral response and even enhancing the antiviral efficiency of IRF1. Collectively, all the data revealed that TRIM21 restricts viral infection in IRF1-dependent and IRF1-independent manners.

## Discussion

As a basic biological process, protein translation is tightly controlled to maintain protein homeostasis. When translation is completed, the nascent proteins are always modified by various manners, such as ubiquitination by TRIM proteins, to confer the specific function, or determine the fate of proteins. The indispensable role of TRIM proteins in post-translational modification has been widely investigated, however, the role of TRIM proteins in regulating protein translation initiation is not clear.

Translation initiation factor eIF2α plays central role in protein translation initiation, which is controlled by phosphorylation of eIF2α by four kinases, PKR, GCN2, PERK and HRI, upon stresses. Among them, PKR is distinct for its role in detection of viral infection and plays essential roles in various biological and physiological processes, such as inflammation, immune response, cancer and aging, all of which are associated with PKR-mediated protein translation (Kroemer G et al., 2022, Derisbourg MJ et al., 2021). PKR is activated by dsRNA produced during viral infection and plays important role in the host immune response. In mammalian cells, at least two signaling pathways are initiated upon detection of cytosolic dsRNA. One is the RLR signaling pathway, and the other is the PKR signaling pathway (Dalet A et al., 2017). A large number of factors, commonly derived from host, have been reported to regulate RLR signaling pathway to achieve a proper and efficient antiviral response (Cao X, 2016, Rehwinkel J and Gack MU, 2020). However, how host factors regulate the stress-activated PKR signaling pathway to balance the protein homeostasis is not clear. It is evidenced that the essential role of TRIM proteins with E3 ligase in regulating RLR signaling pathway (van Wijk SJ et al., 2019), the role of TRIM proteins in PRR signaling pathways and immune-related diseases, while the function of TRIM-mediated ubiquitination in PKR signaling is not well studied. In present study, we firstly disclose that the E3 ligase TRIM21 inhibits protein translation initiation by functioning on PKR signaling pathway upon stresses, such as viral infection and TG treatment, providing new insight into the regulation of protein synthesis by TRIM proteins.

TRIM21 is a member of the TRIM superfamily, which has been reported to be involved in diverse biological processes and implicated in various diseases (Wang L and Ning S, 2021). Our previous study showed an important role of TRIM21 in the virus-triggered RLR signaling pathway (Lee AYS et al., 2021). Whether TRIM21 regulates the PKR signaling pathway remains elusive. In the present study, we find that TRIM21 negatively regulates PKR activation by promoting K6-linked ubiquitination of PP1α under stress. Several lines of evidence strongly support our conclusion. First, knocking out *TRIM21* augments stress-induced PKR activation and subsequent inhibition of protein synthesis. Second, although TRIM21 has no effect on PKR ubiquitination, the E3 ligase activity is essential for TRIM21-mediated inactivation of PKR, suggesting an indirect regulation of PKR by TRIM21. Third, TRIM21 specifically abolishes PKR phosphorylation but not dsRNA detection or dimerization of PKR, indicating that the phosphatases of PKR may be regulated by TRIM21. Fourth, PP1α, the phosphatase of PKR, is responsible for TRIM21-mediated inhibition of PKR activation. TRIM21 promotes its interaction with PP1α and catalyzes the K6-linked ubiquitination of PP1α on Lys60 under stress. Fifth, silencing PP1α or K60 mutation of PP1α resulting in its inactivation abolishes the inhibition of PKR phosphorylation by TRIM21. Collectively, the TRIM21-PP1α axis is defined as a newly discovered program for regulating stress-induced PKR-mediated protein synthesis. Moreover, we find an IFN-independent antiviral function of TRIM21 by reversing PKR-mediated inhibition of the protein synthesis of antiviral effectors. TRIM21 constitutively inhibits the replication of VSV or SeV in IFNAR1-deficient cells. We identify several previously known and unknown antiviral effectors regulated by the TRIM21-PKR axis, which broadens our understanding of antiviral genes and strengthens the evidence that the antiviral function of TRIM21 is achieved by reversing PKR-mediated translational shutdown.

As one of the classic posttranslational modifications (PTMs), ubiquitination profoundly affects the fundamental physiological processes, such as the cell proliferation, differentiation, cell death, and the protein stability and structure, of all species (van Wijk SJ et al., 2019, Berndsen CE and Wolberger C, 2014). E3 ligases are vital components in this process, directly interacting with and catalyzing different ubiquitin linkages of their specific substrates, which can occur through K6-, K11-, K27-, K29-, K33-, K48- and K63-linked ubiquitination (Cruz Walma DA et al., 2022). Several types of ubiquitin linkages catalyzed by TRIM21 have been reported, and different ubiquitin linkages play distinct roles in host defense against pathogenic invasion (Jones EL et al., 2021). For instance, TRIM21-mediated K48-linked ubiquitination is tightly related to proteasome-dependent degradation, which results in the degradation of viral proteins in virus-infected cells (Mu T et al., 2020, Song Y et al., 2021, Hauler F et al., 2012). However, K48-linked ubiquitination of DDX41, one of the sensors detecting dsDNA, results in the inhibition of the innate immune response triggered by DNA virus (Zhang Z et al., 2011). K63-linked ubiquitination catalyzed by TRIM21 augments the activation of the intracellular antibody-triggered innate immune response (McEwan WA et al., 2013). K27-linked ubiquitination promotes RNA virus-induced MAVS activation and the innate immune response to viral infection (Xue B et al., 2018). Importantly, in this study, we find that TRIM21-catalyzed K6-linked ubiquitination of PP1α restricts viral infection by promoting the translation of intrinsic antiviral effectors. Thus, our findings indicate a newly discovered type of ubiquitin linkage mediated by TRIM21 that can participate in the host antiviral response with a distinct mechanism.

Virus-induced PKR activation suppresses the synthesis of interferon-stimulated genes (ISGs) upon viral infection (Feng H et al., 2017, Garaigorta U and Chisari FV, 2009). Emerging evidence demonstrates the negative role of PKR-triggered protein inhibition in the host defense against viral infection. For example, one recent study reported that NLRX1 is a positive regulator in the antiviral response by limiting PKR-mediated translational shutoff of the IRF1 protein, which results in inhibition production of IRF1-dependent antiviral factors (Darini C et al., 2019). Similarly, our data show an IRF1-dependent antiviral efficiency regulated by TRIM21. TRIM21 promotes IRF1 protein synthesis by inhibiting PKR activation upon virus infection. Combination of the positive role of TRIM21 in innate immunity, we define the antiviral role of TRIM21 both in augmenting the transcription of IFN production and in facilitating the translation of IRF1 protein. Besides innate antiviral response, IRF1 appreciates its adaptive immunity and tumor suppression. Given that, we speculate that TRIM21 may have a role, not limited by viral infection, but also tumorigenesis, in adaptive immunity, which needs further study. In addition, we also find an IRF1-independent antiviral function mediated by TRIM21 via analyzing the proteomics. Some previously unknown antiviral factors have been discovered, all of which were directly regulated by TRIM21. Together with these findings, we demonstrate that an IFN-independent antiviral role of TRIM21 restricts viral infection by reversing PKR-mediated protein translational inhibition of the antiviral factors. Collectively, the results of our study highlight a newly discovered biological role of TRIM21, and these data provide new evidence of PKR-mediated translational arrest in host resistance to virus infection.

## Materials and methods

### Ethics Statement

All animal experiments were carried out under the supervision of the Institution Animal Care and Use Committee of Hunan University.

### Mice

C57BL/6 mice (6-8 weeks of age) were purchased from Hunan SJA Laboratory Animal Co. Ltd. and kept under specific pathogen-free conditions in the animal care facility of Hunan University. All animal experiments were conducted in accordance with the guidelines of the Laboratory Animal Management Regulation with approval of Hunan University. To knock down the expression of *Trim21* in mouse tissues, lentiviruses expressing specific shRNAs were produced and purified. Weight- and sex-matched mice were arbitrarily assigned to two groups: one group was injected with sh-vector (1x10^11^ PFU/g), and the other group was injected with sh-*Trim21* (1x10^11^ PFU/g). All the animals received an intravenous injection in the tail once every three days for a total of five times. For in vivo viral infection, weight- and sex-matched mice were injected with VSV intravenously (1x10^8^ PFU/g) for 36 h. The livers, spleens and lungs were isolated to determine the VSV RNA levels and the protein level of Trim21.

### Cell culture and reagents

The HLCZ01 cell line, a hepatoma cell line supporting the entire life cycle of HCV and HBV, was previously established in our laboratory. HEK293T cells were purchased from Boster, Huh7.5 cells were kindly provided by Charles M. Rice (Rockefeller University, New York), and Huh7.5 and A549 cells were obtained from the American Type Culture Collection. HLCZ01 cells were cultured in collagen-coated tissue culture plates containing Dulbecco’s modified Eagle medium (DMEM)–F-12 medium supplemented with 10% (vol/vol) fetal bovine serum (FBS) (Gibco), 40 ng/ml dexamethasone (Sigma), insulin-transferrin-selenium (ITS) (Lonza), and penicillin–streptomycin (Thermo Fisher Scientific). Other cells were propagated in DMEM supplemented with 10% FBS, nonessential amino acid solution (Thermo Fisher Scientific), and penicillin–streptomycin. Cell transfection with plasmids was conducted using ViaFect™ Transfection Reagent (Promega) or Lipofectamine® 2000 (Thermo Fisher Scientific) in Opti-MEMTM medium (Thermo Fisher Scientific). For stable transfection or infection, monoclonal cells were screened using the antibiotic Puro (Gibco).

### Virus

The pJFH1 plasmid was a gift from Takaji Wakita (National Institute of Infectious Diseases, Tokyo, Japan). The linearized DNA from the pJFH1 plasmid was purified and used as the template for in vitro transcription with a MEGAscript kit (Ambion, Austin, TX). In vitro-transcribed genomic JFH1 RNA was delivered into Huh7.5 cells by electroporation. The transfected cells were cultured for the indicated periods. The cells were passaged every 3 to 5 days, while the corresponding supernatants were collected and filtered with a 0.45 μm filter device. The viral titers are presented as focus-forming units (FFUs) per milliliter, determined as the average number of NS5A-positive foci detected in Huh7.5 cells. SeV and VSV were kindly shared by Xuetao Cao (Second Military Medical University, Shanghai, China).

### Antibodies

Antibodies against the following proteins were obtained from commercial sources and used for immunoblots: anti-flag-tag (F3165, Sigma), anti-V5-tag (R960-25, Thermo Fisher Scientific), anti-puromycin (MABE343, Merck Millipore), anti-GAPDH (MAB374, Merck Millipore), anti-phospho-eIF2α Ser51 (3398, CST), anti-TRIM21 (92043, CST), anti-PKR (12297, CST), anti-Myc tag (2276, CST), anti-HA tag (ab236632, Abcam), anti-TRIM21 (ab91423, Abcam), anti-phospho-PKR T451 (ab81303, Abcam), and anti-IRF1 (8478, CST). Anti-interferon alpha/beta receptor 1 (ab45172, Abcam), anti-STAT2 (72604, CST), anti-PP1α (ab137512, Abcam), anti-ISG15 (ab285367, Abcam), anti-GST mouse monoclonal antibody (HT601-01, Transgen), β-actin (A5541, Sigma), and goat anti-mouse IgG (HRP-linked) (AP124P, Merck Millipore) were also used. The following antibodies were obtained from commercial sources and used for immunofluorescence staining: anti-PKR (sc6282, Santa Cruz Biotechnology, USA), anti-PP1α (ab137512, Abcam) and donkey anti-rabbit IgG (H + L) highly cross-adsorbed secondary antibody conjugated with Alexa Fluor 594 (A-21207, Thermo Fisher Scientific). The mouse monoclonal anti-HCV core antibody was a gift from Chen Liu.

### Plasmid construction

TRIM21, PP1α, and PKR cDNAs were synthesized from total cellular RNA isolated from HLCZ01 cells by standard reverse transcription-PCR (RT–PCR). Subsequently, they were cloned into the pcDNA3.1a vector, p3FLAG-CMV vector pCMV-N-Myc or pGEX4T2 vector. Multiple domains of TRIM21 and PP1α were amplified from the templates of full-length TRIM21 and PP1α, which were then cloned into p3xFLAG-CMV. The primers for amplifying these genes are listed in Table 1. The pHA-Ub (K6, K11, K27, K29 and K33) plasmids were kindly shared by Hongbing Shu (Wuhan University). The pHA-Ub (WT, K48 and K63) plasmids were kindly provided by Zhengfan Jiang (Peking University).

**Table 1.**
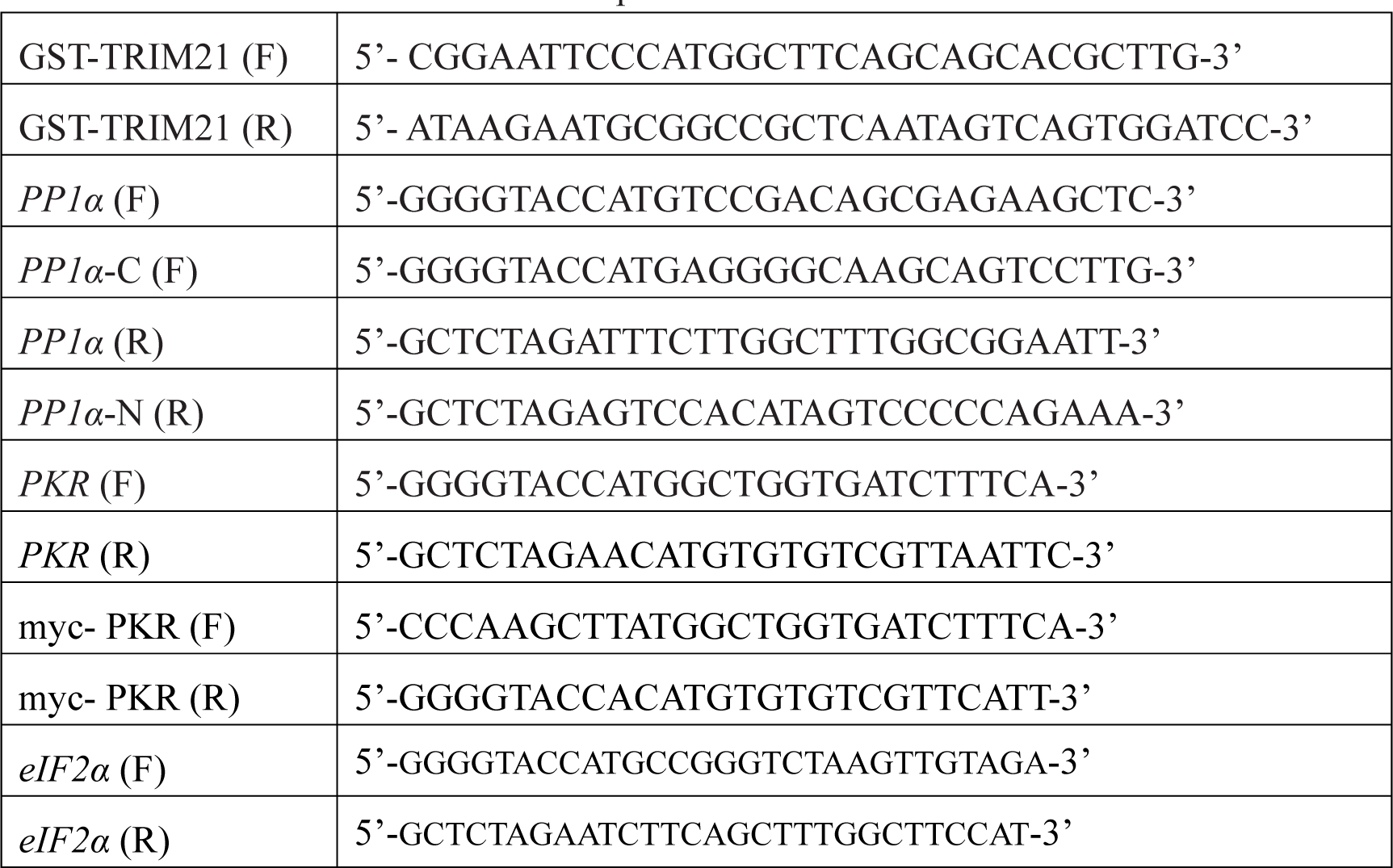
Primers for the construction of plasmids.

### Lentivirus production and generation of stable cell lines

HEK293T cells plated in 10 cm dishes were transfected with 8 μg packaging plasmid psPAX2, 2.7 μg envelope plasmid pMD2G and 8 μg target plasmid encoding shRNA or lentiCRISPRv2 encoding sgRNA using Lipofectamine® 2000 (Thermo Fisher Scientific). Lentivirus supernatants were collected at 36 h, 48 h, 56 h and 72 h posttransfection and clarified by filtration through 0.45 μm syringe filters. HLCZ01, A549 or Huh7.5 cells were plated in 6-well plates prior to being transduced with 500 μL per well of lentivirus supernatant. Transduced cells were selected for by the addition of 2 μg/mL puromycin at 72 h post-transduction. Loss of target protein expression was confirmed by Western blot. The target sequences used for shRNA gene silencing and lentiCRISPRv2 encoding sgRNA plasmids are listed in Table 2.

**Table 2.**
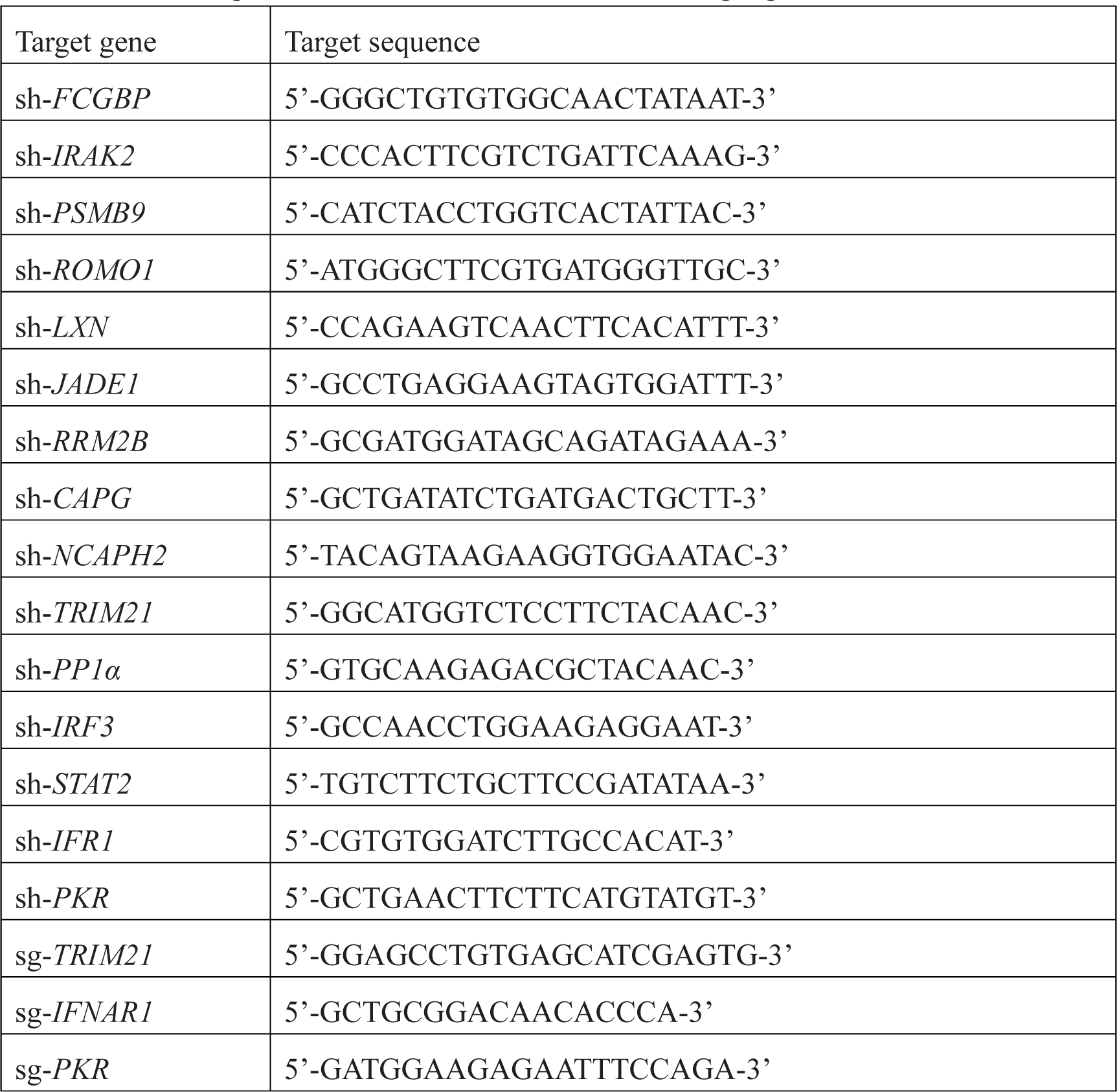
Short hairpin RNAs used for knockdowns and single guide RNAs used for CRISPR/Cas9.

### Real-time PCR assay

Total cellular RNA was extracted by TRIzol reagent (Invitrogen) according to the manufacturer’s protocol. The Superscript III first-strand synthesis kit for reverse transcription of RNA was purchased from Invitrogen. After RQ1 DNase (Promega) treatment, the extracted RNA was used as the template for reverse transcription-PCR. Real-time PCR was performed as described previously (51). GAPDH was used as the internal control. The primers used for real-time PCR are listed in Table 3.

**Table 3.**
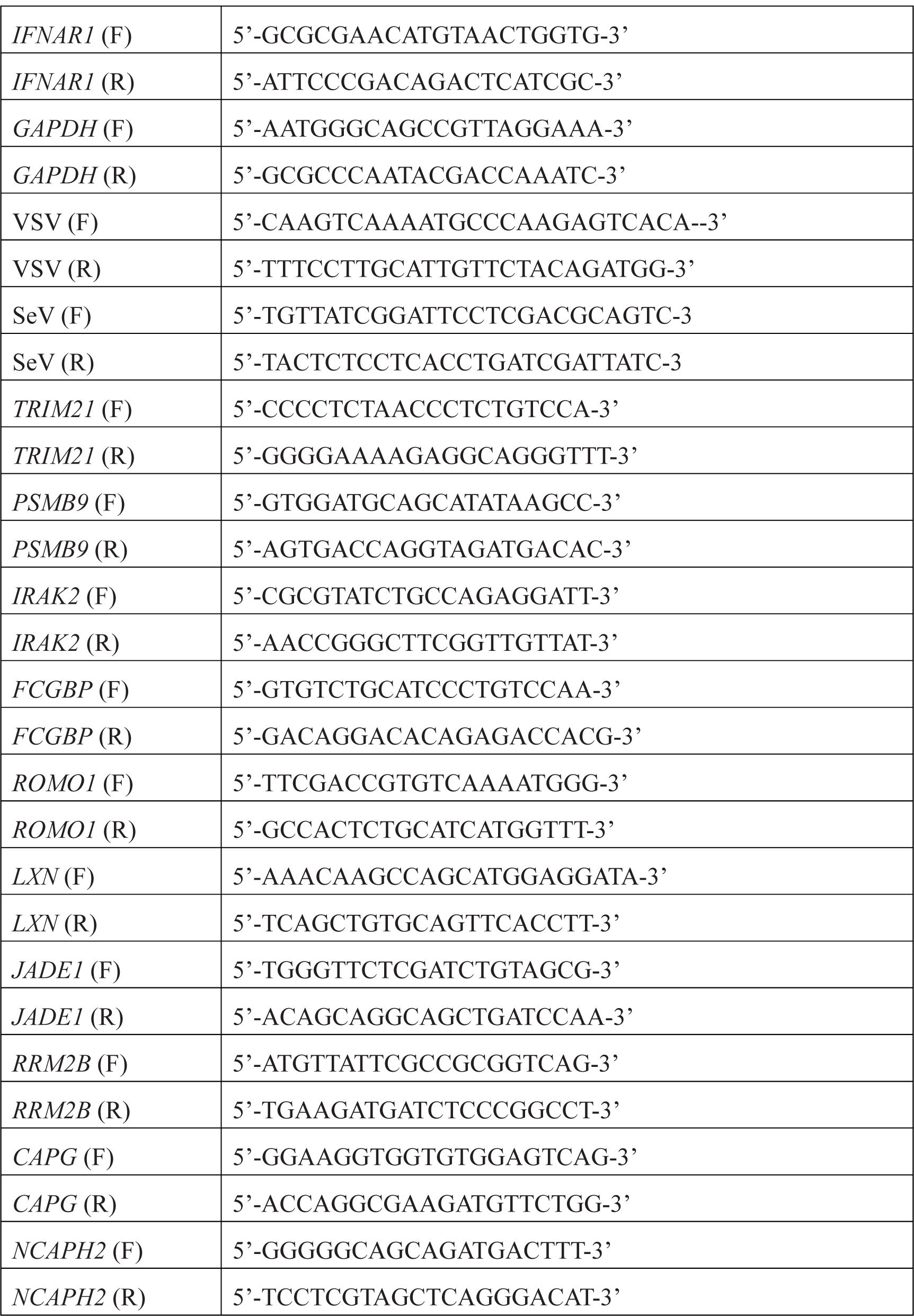
Primers for Real-time PCR.

### Western blotting

Cells were washed with PBS and lysed with RIPA buffer (Thermo Fisher Scientific) supplemented with protease inhibitor cocktail (Thermo Fisher Scientific). The lysates were incubated on ice for 30 min and centrifuged at 16000 g for 15 min at 4 °C. Proteins were resolved on SDS–PAGE gels and transferred to polyvinylidene difluoride (PVDF) membranes (Merck Millipore). The PVDF membranes were then blocked with 5% skim milk and sequentially incubated with primary and secondary antibodies. The bound antibodies were detected using SuperSignal West Pico chemiluminescent substrate (Pierce, Rockford, IL).

### IP and immunoblotting

Cells were washed with PBS and lysed with Pierce™ IP Lysis Buffer (Thermo Fisher Scientific) supplemented with protease inhibitor cocktail. The lysates were incubated on ice for 30 min and centrifuged at 12000 g for 15 min at 4 °C. The lysates were diluted to a concentration of 2 μg/μL with PBS before IP. The lysates (200 μg) were immunoprecipitated with the indicated antibodies. The immunocomplex was captured by adding Protein G Agarose (Merck Millipore, Darmstadt, Germany). The protein binding to the beads was boiled in 2 x Laemmli sample buffer (Bio–Rad, Hercules) and was then subjected to SDS–PAGE.

### RNA immunoprecipitation

The cells were trypsinized to detach them, and the supernatant was discarded. The cells were washed by gently resuspending them in 1 ml PBS and pelleted by centrifugation at 3,000 *g* for 1 min. Then, the cells were resuspended in 1 mL 1% formaldehyde (diluted in PBS) and allowed to stand for 10 min at room temperature. The cells were pelleted by centrifugation at 3,000 *g* for 1 min, resuspended in 1 mL 0.25 M glycine solution (diluted in PBS) and again allowed to stand for 10 min at room temperature. The cells were pelleted and washed with 500 μL PBS, and then the cells were lysed with RIPA buffer supplemented with a protease inhibitor cocktail and an RNase inhibitor on ice for 30 min. The subsequent immunoprecipitate was processed as described above. Protein–RNA complexes binding to beads were eluted in PBS at 70 °C for 45 min. The eluted material was lysed in ice-cold TRIzol reagent for RT–PCR.

### Immunofluorescence staining

Cells were seeded into a confocal dish and fixed with 4% paraformaldehyde for 15 min at room temperature. The cells were washed with PBS, permeabilized for 15 min with 0.2% Triton X-100 in PBS for 10 min, blocked with goat serum (Thermo Fisher Scientific) for 30 min at room temperature, and sequentially incubated with primary and fluorescence-labeled secondary antibodies (Invitrogen) (diluted in PBS to 1:500) at room temperature for 2 h. Nuclei were counterstained with DAPI (Vector Laboratories, Burlingame) for 5 min. Images were captured using a TI-E + A1 SI confocal microscope (Nikon).

### Plaque assay

The supernatants were diluted into multiple concentrate gradients with medium without FBS and transduced into VERO cells. After 1 h, the cells were placed in complete medium. Two hours later, 1% agarose media was added on top of the virus-infected Vero cells. After 1 day, cells with plaques were fixed with 4% paraformaldehyde for 30 min at room temperature and stained with 0.1% crystal violet.

### Puromycin incorporation assay

Cells were pulsed with puromycin (10 µg/mL) for 1 h and then fixed with 4% paraformaldehyde, followed by staining with anti-puromycin and DAPI. Cells were imaged by confocal microscopy. Alternatively, cells were pulsed with puromycin (10 µg/mL) for 1 h, lysed, and puromycin-labeled proteins were identified by immunoblot analysis.

### Statistical Analysis

All results are presented as the means and standard deviations (SDs). Comparisons between two groups were performed by using Student’s *t* test.

## Acknowledgments

We thank Charles M. Rice for the Huh7.5 cells, Takaji Wakita (National Institute of Infectious Diseases, Tokyo, Japan) for pJFH1 plasmid, Stanley Lemon (University of North Carolina, Chapel Hill, NC) for pH 77-S plasmid, Jianguo Wu (Jinan University, Guangzhhou, China) for VSV, and Xingyi Ge (Hunan University, Changsha, China) for SeV. We also thank Chen Liu (Yale University, New Haven, USA), Hongbing Shu (Wuhan University, Wuhan, China) and Zhengfan Jiang (Peking University, Beijing, China) for kindly sharing research materials. This work was Supported by grants from the National Natural Science Foundation of China (81730064, 82072269, 81902069 and 81571985), National Major Science and Technology Projects of China (2017ZX10202201-005 and 2009ZX10004-312), Postdoctoral Research Foundation of China (2019M652760) and Hunan Provincial Innovation Foundation For Postgraduate (CX20210395)

## Author Contributions

L, H., X, B. and Z, H. designed the experiments and wrote the manuscript. L, H., X, B., L, S., F, Q. W, M and L, X performed the experiments. L, H., X, B., D, R., X, Y., W, J., W, X., T, R., C, S., W, L and L, Q analyzed the data. P, Y analyzed the data. T, S., X, B. and Z, H. provided the research materials. X, B. and Z, H. supervised the experiments.

## References

1. Berndsen CE, Wolberger C (2014) New insights into ubiquitin E3 ligase mechanism. Nat Struct Mol Biol 21: 301–7

2. Cao X (2016) Self-regulation and cross-regulation of pattern-recognition receptor signalling in health and disease. Nat Rev Immunol 16: 35–50

3. Chen R, Ishak CA, De Carvalho DD (2021) Endogenous Retroelements and the Viral Mimicry Response in Cancer Therapy and Cellular Homeostasis. Cancer Discov 11: 2707–2725

4. Chen YG, Hur S (2022) Cellular origins of dsRNA, their recognition and consequences. Nat Rev Mol Cell Biol 23: 286–301

5. Chevrier N, Mertins P, Artyomov MN, Shalek AK, Iannacone M, Ciaccio MF, Gat-Viks I, Tonti E, DeGrace MM, Clauser KR, Garber M, Eisenhaure TM, Yosef N, Robinson J, Sutton A, Andersen MS, Root DE, von Andrian U, Jones RB, Park H et al. (2011) Systematic discovery of TLR signaling components delineates viral-sensing circuits. Cell 147: 853–67

6. Cronk JC, Herz J, Kim TS, Louveau A, Moser EK, Sharma AK, Smirnov I, Tung KS, Braciale TJ, Kipnis J (2017) Influenza A induces dysfunctional immunity and death in MeCP2-overexpressing mice. JCI Insight 2: e88257

7. Cruz Walma DA, Chen Z, Bullock AN, Yamada KM (2022) Ubiquitin ligases: guardians of mammalian development. Nat Rev Mol Cell Biol

8. Dalet A, Arguello RJ, Combes A, Spinelli L, Jaeger S, Fallet M, Vu Manh TP, Mendes A, Perego J, Reverendo M, Camosseto V, Dalod M, Weil T, Santos MA, Gatti E, Pierre P (2017) Protein synthesis inhibition and GADD34 control IFN-beta heterogeneous expression in response to dsRNA. EMBO J 36: 761–782

9. Dalet A, Gatti E, Pierre P (2015) Integration of PKR-dependent translation inhibition with innate immunity is required for a coordinated anti-viral response. FEBS Lett 589: 1539–1545

10. Darini C, Ghaddar N, Chabot C, Assaker G, Sabri S, Wang S, Krishnamoorthy J, Buchanan M, Aguilar-Mahecha A, Abdulkarim B, Deschenes J, Torres J, Ursini-Siegel J, Basik M, Koromilas AE (2019) An integrated stress response via PKR suppresses HER2+ cancers and improves trastuzumab therapy. Nat Commun 10: 2139

11. Dauber B, Wolff T (2009) Activation of the Antiviral Kinase PKR and Viral Countermeasures. Viruses 1: 523–544

12. Derisbourg MJ, Hartman MD, Denzel MS (2021) Perspective: Modulating the integrated stress response to slow aging and ameliorate age-related pathology. Nat Aging 1: 760–768

13. Doyle T, Moncorge O, Bonaventure B, Pollpeter D, Lussignol M, Tauziet M, Apolonia L, Catanese MT, Goujon C, Malim MH (2018) The interferon-inducible isoform of NCOA7 inhibits endosome-mediated viral entry. Nat Microbiol 3: 1369–1376

14. Feng H, Lenarcic EM, Yamane D, Wauthier E, Mo J, Guo H, McGivern DR, Gonzalez-Lopez O, Misumi I, Reid LM, Whitmire JK, Ting JP, Duncan JA, Moorman NJ, Lemon SM (2017) NLRX1 promotes immediate IRF1-directed antiviral responses by limiting dsRNA-activated translational inhibition mediated by PKR. Nat Immunol 18: 1299–1309

15. Galanina N, Goodman AM, Cohen PR, Frampton GM, Kurzrock R (2018) Successful Treatment of HIV-Associated Kaposi Sarcoma with Immune Checkpoint Blockade. Cancer Immunol Res 6: 1129–1135

16. Garaigorta U, Chisari FV (2009) Hepatitis C virus blocks interferon effector function by inducing protein kinase R phosphorylation. Cell Host Microbe 6: 513–522

17. Hauler F, Mallery DL, McEwan WA, Bidgood SR, James LC (2012) AAA ATPase p97/VCP is essential for TRIM21-mediated virus neutralization. Proc Natl Acad Sci U S A 109: 19733–8

18. Hotamisligil GS (2010) Endoplasmic reticulum stress and the inflammatory basis of metabolic disease. Cell 140: 900–917

19. Hsu LC, Park JM, Zhang K, Luo JL, Maeda S, Kaufman RJ, Eckmann L, Guiney DG, Karin M (2004) The protein kinase PKR is required for macrophage apoptosis after activation of Toll-like receptor 4. Nature 428: 341–5

20. Jha HC, A JM, Saha A, Banerjee S, Lu J, Robertson ES (2014) Epstein-Barr virus essential antigen EBNA3C attenuates H2AX expression. J Virol 88: 3776–3788

21. Jones EL, Laidlaw SM, Dustin LB (2021) TRIM21/Ro52 - Roles in Innate Immunity and Autoimmune Disease. Front Immunol 12: 738473

22. Koga R, Kubota M, Hashiguchi T, Yanagi Y, Ohno S (2018) Annexin A2 Mediates the Localization of Measles Virus Matrix Protein at the Plasma Membrane. J Virol 92: e00181–18

23. Kroemer G, Galassi C, Zitvogel L, Galluzzi L (2022) Immunogenic cell stress and death. Nat Immunol

24. Lee AYS, Reed JH, Gordon TP (2021) Anti-Ro60 and anti-Ro52/TRIM21: Two distinct autoantibodies in systemic autoimmune diseases. J Autoimmun 124: 102724

25. Liu G, Gack MU (2020) Distinct and Orchestrated Functions of RNA Sensors in Innate Immunity. Immunity 53: 26–42

26. Liu S, Cai X, Wu J, Cong Q, Chen X, Li T, Du F, Ren J, Wu YT, Grishin NV, Chen ZJ (2015) Phosphorylation of innate immune adaptor proteins MAVS, STING, and TRIF induces IRF3 activation. Science 347: aaa2630

27. Liuyu T, Yu K, Ye L, Zhang Z, Zhang M, Ren Y, Cai Z, Zhu Q, Lin D, Zhong B (2019) Induction of OTUD4 by viral infection promotes antiviral responses through deubiquitinating and stabilizing MAVS. Cell Res 29: 67–79

28. Lu B, Nakamura T, Inouye K, Li J, Tang Y, Lundback P, Valdes-Ferrer SI, Olofsson PS, Kalb T, Roth J, Zou Y, Erlandsson-Harris H, Yang H, Ting JP, Wang H, Andersson U, Antoine DJ, Chavan SS, Hotamisligil GS, Tracey KJ (2012) Novel role of PKR in inflammasome activation and HMGB1 release. Nature 488: 670–674

29. Lu X, Chen Q, Liu H, Zhang X (2021) Interplay Between Non-Canonical NF-kappaB Signaling and Hepatitis B Virus Infection. Front Immunol 12: 730684

30. McCormick C, Khaperskyy DA (2017) Translation inhibition and stress granules in the antiviral immune response. Nat Rev Immunol 17: 647–660

31. McEwan WA, Tam JC, Watkinson RE, Bidgood SR, Mallery DL, James LC (2013) Intracellular antibody-bound pathogens stimulate immune signaling via the Fc receptor TRIM21. Nat Immunol 14: 327–36

32. Mu T, Zhao X, Zhu Y, Fan H, Tang H (2020) The E3 Ubiquitin Ligase TRIM21 Promotes HBV DNA Polymerase Degradation. Viruses 12. 346

33. Pakos-Zebrucka K, Koryga I, Mnich K, Ljujic M, Samali A, Gorman AM (2016) The integrated stress response. EMBO Rep 17: 1374–1395

34. Qiao H, Jiang T, Mu P, Chen X, Wen X, Hu Z, Tang S, Wen J, Deng Y (2021) Cell fate determined by the activation balance between PKR and SPHK1. Cell Death Differ 28: 401–418

35. Rehwinkel J, Gack MU (2020) RIG-I-like receptors: their regulation and roles in RNA sensing. Nat Rev Immunol 20: 537–551

36. Sharma NR, Majerciak V, Kruhlak MJ, Zheng ZM (2017) KSHV inhibits stress granule formation by viral ORF57 blocking PKR activation. PLoS Pathog 13: e1006677

37. Song Y, Li M, Wang Y, Zhang H, Wei L, Xu W (2021) E3 ubiquitin ligase TRIM21 restricts hepatitis B virus replication by targeting HBx for proteasomal degradation. Antiviral Res 192: 105107

38. Talloczy Z, Jiang W, Virgin HWt, Leib DA, Scheuner D, Kaufman RJ, Eskelinen EL, Levine B (2002) Regulation of starvation- and virus-induced autophagy by the eIF2alpha kinase signaling pathway. Proc Natl Acad Sci U S A 99: 190–5

39. Tan SL, Tareen SU, Melville MW, Blakely CM, Katze MG (2002) The direct binding of the catalytic subunit of protein phosphatase 1 to the PKR protein kinase is necessary but not sufficient for inactivation and disruption of enzyme dimer formation. J Biol Chem 277: 36109–36117

40. Tatematsu M, Nishikawa F, Seya T, Matsumoto M (2013) Toll-like receptor 3 recognizes incomplete stem structures in single-stranded viral RNA. Nat Commun 4: 1833

41. van Wijk SJ, Fulda S, Dikic I, Heilemann M (2019) Visualizing ubiquitination in mammalian cells. EMBO Rep 20: e46520

42. Wang J, Li H, Xue B, Deng R, Huang X, Xu Y, Chen S, Tian R, Wang X, Xun Z, Sang M, Zhu H (2020) IRF1 Promotes the Innate Immune Response to Viral Infection by Enhancing the Activation of IRF3. J Virol 94: e01231–20

43. Wang K, Lai C, Li T, Wang C, Wang W, Ni B, Bai C, Zhang S, Han L, Gu H, Zhao Z, Duan Y, Yang X, Xing L, Zhao L, Zhou S, Xia M, Jiang C, Wang X, Yang P (2018) Basic fibroblast growth factor protects against influenza A virus-induced acute lung injury by recruiting neutrophils. J Mol Cell Biol 10: 573–585

44. Wang L, Ning S (2021) TRIMming Type I Interferon-Mediated Innate Immune Response in Antiviral and Antitumor Defense. Viruses 13: 279

45. Wang W, Jin Y, Zeng N, Ruan Q, Qian F (2017) SOD2 Facilitates the Antiviral Innate Immune Response by Scavenging Reactive Oxygen Species. Viral Immunol 30: 582–589

46. Wek RC (2018) Role of eIF2alpha Kinases in Translational Control and Adaptation to Cellular Stress. Cold Spring Harb Perspect Biol 10: a032870

47. Wu J, Chen ZJ (2014) Innate immune sensing and signaling of cytosolic nucleic acids. Annu Rev Immunol 32: 461–488

48. Xue B, Li H, Guo M, Wang J, Xu Y, Zou X, Deng R, Li G, Zhu H (2018) TRIM21 Promotes Innate Immune Response to RNA Viral Infection through Lys27-Linked Polyubiquitination of MAVS. J Virol 92: e00321–18

49. Yamane D, Feng H, Rivera-Serrano EE, Selitsky SR, Hirai-Yuki A, Das A, McKnight KL, Misumi I, Hensley L, Lovell W, Gonzalez-Lopez O, Suzuki R, Matsuda M, Nakanishi H, Ohto-Nakanishi T, Hishiki T, Wauthier E, Oikawa T, Morita K, Reid LM et al. (2019) Basal expression of interferon regulatory factor 1 drives intrinsic hepatocyte resistance to multiple RNA viruses. Nat Microbiol 4: 1096–1104

50. Yang D, Zuo C, Wang X, Meng X, Xue B, Liu N, Yu R, Qin Y, Gao Y, Wang Q, Hu J, Wang L, Zhou Z, Liu B, Tan D, Guan Y, Zhu H (2014) Complete replication of hepatitis B virus and hepatitis C virus in a newly developed hepatoma cell line. Proc Natl Acad Sci U S A 111: E1264–1273

51. Yang W, Wu YH, Liu SQ, Sheng ZY, Zhen ZD, Gao RQ, Cui XY, Fan DY, Qin ZH, Zheng AH, Wang PG, An J (2020) S100A4+ macrophages facilitate zika virus invasion and persistence in the seminiferous tubules via interferon-gamma mediation. PLoS Pathog 16: e1009019

52. Zhang T, Yin C, Fedorov A, Qiao L, Bao H, Beknazarov N, Wang S, Gautam A, Williams RM, Crawford JC, Peri S, Studitsky V, Beg AA, Thomas PG, Walkley C, Xu Y, Poptsova M, Herbert A, Balachandran S (2022) ADAR1 masks the cancer immunotherapeutic promise of ZBP1-driven necroptosis. Nature

53. Zhang Z, Yuan B, Bao M, Lu N, Kim T, Liu YJ (2011) The helicase DDX41 senses intracellular DNA mediated by the adaptor STING in dendritic cells. Nat Immunol 12: 959–965

